# Molecular glue degraders enhance CAPRIN1-dependent lysosomal degradation of APP in Alzheimer’s disease

**DOI:** 10.1101/2023.12.29.573648

**Authors:** Sunghan Jung, Raktim Roy, Xu Wang, Bin Liu, Nancy Jaiswal, Ratan K. Rai, Yuichiro Takagi, Cen Gao, Lifan Zeng, Ho-Yin Lo, Reagan K. Wohlford, Ryan K. Higgins, Anantha Shekhar, Anita C. Bellail, Chunhai Hao

## Abstract

Overexpression of amyloid precursor protein (APP) is a key driver of amyloid β (Aβ) pathology, making APP a compelling therapeutic target in Alzheimer’s disease (AD). Here, we identify small molecules that selectively degrade APP in human neurons through a targeted protein degradation approach. Structural analyses reveal that these compounds act as molecular glues, binding at the interface between cytoplasmic activation/proliferation-associate protein 1 (CAPRIN1) and APP and stabilizing their interaction within a ternary complex. In neurons derived from induced pluripotent stem cells (iPSCs) from sporadic and familial AD patients, these compounds promote CAPRIN1-dependent degradation of both wild-type and mutant APP through the endo-lysosomal pathway. The lead compound, 0152, is blood-brain barrier permeable, and its administration significantly reduces Aβ production and amyloid burden in the brains of 5xFAD mice. These findings demonstrate that molecular glue degraders harness the CAPRIN1-dependent lysosomal pathway for selective APP degradation, offering a targeted therapeutic strategy for AD.

## INTRODUCTION

Alzheimer’s disease (AD) is a progressive neurodegenerative disorder and the leading cause of dementia worldwide, for which curative treatments remain elusive^1–3^. Pathologically, AD is characterized by the extracellular accumulation of amyloid-β (Aβ) peptides into plaques and the intracellular aggregation of hyperphosphorylated tau into neurofibrillary tangles, predominantly within the cerebral cortex and hippocampus^4^. According to the amyloid cascade hypothesis, the overproduction and aggregation of Aβ peptides act as the initial event that drives downstream tau pathology, neuroinflammation, and neuronal loss^5, 6^. Aβ peptides are generated via the sequential cleavage of amyloid precursor protein (APP) by β-secretase 1 (BACE1) and γ-secretase in neurons, followed by their extracellular release and aggregation into plaques. Therapeutic strategies aimed at interrupting this cascade have led to the development of several disease-modifying agents, including BACE1 and γ-secretase inhibitors, as well as monoclonal antibodies (mAbs) targeting Aβ^6^. While secretase inhibitors have failed to show clinical efficacy, mAbs such as aducanumab, lecanemab, and donanemab have received FDA approval for patients with early-stage AD and mild dementia^7–9^, underscoring the therapeutic potential of targeting APP-derived Aβ^10^.

The *APP* gene and its expression levels critically influence the onset and progression of AD and related dementias (ADRD). In Down syndrome (DS), trisomy 21 leads to APP gene triplication, resulting in elevated APP expression and Aβ production, which drive AD-like neuropathology that typically appears before age 40^11^. Similarly, early-onset familial AD (fAD), which manifests before age 65, is driven by mutations in *APP*, *Presenilin-1* (*PSEN1*) or *PSEN2* that promote increased APP expression and generation of aggregation-prone Aβ42 peptides^12^. Late-onset sporadic AD (sAD), which arises after age 60, is multifactorial and involves mechanisms such as neuronal genomic mosaicism and RNA dysregulation, both contributing to elevated APP expression and Aβ accumulation^13^. Aging remains the strongest risk factor for neurodegeneration^14^. Notably, recent findings have shown increased APP and Aβ levels in the cortical and hippocampal neurons of cognitively impaired octogenarians, while genetic ablation of the *appa* gene in the turquoise killifish ameliorates age-related cognitive decline^15^. Together, these observations highlight APP as a compelling therapeutic target not only for AD and ADRD but also for age-associated cognitive decline.

Targeted protein degradation (TPD) has emerged as a powerful drug discovery strategy that employs small molecules to selectively eliminate disease-associated proteins via proteasomal or lysosomal pathways^16, 17^. Most degraders developed to date are large, bifunctional molecules such as proteolysis-targeting chimeras (PROTACs) and lysosome-targeting chimeras (LYTACs) that have primarily been applied to oncogenic proteins^18, 19^. Some PROTACs have been applied to neurodegenerative targets including tau^20^. In contrast, monovalent molecular glue degraders represent a mechanistically distinct class of small molecules that bind at the interface between an ubiquitin E3 ligase and a substrate protein, inducing de novo protein–protein interactions or stabilizing weak pre-existing ones to promote E3 ligase-mediated proteasomal degradation^21–23^. However, the discovery of such degraders has been limited to the small number of E3 ligases amenable to this mechanism^24^. Here, we report a new class of molecular glue degrader that act at the interface between cytoplasmic activation/proliferation-associate protein 1 (CAPRIN1) and APP. These compounds enhance the CAPRIN1–APP interaction, facilitating CAPRIN1-mediated lysosome degradation of APP in neurons derived from induced pluripotent stem cells (iPSCs) from AD patients. This approach significantly reduces Aβ42 plaque deposition in AD mouse models and offers a promising therapeutic avenue for the treatment of AD.

## RESULTS

### Discovery of active compounds as APP degraders through cell-based screening

We previously reported CAPRIN1-targeted molecular degraders that bind to CAPRIN1 and F-box protein 42 (FBXO42), promoting FBXO42-mediated proteasomal degradation of small ubiquitin-related modifier 1 (SUMO1)^25^ in cancers^26^. Motivated by proteomic evidence of the interaction between CAPRIN1 and APP^27^, we investigated whether CAPRIN1-targeted molecules could also promote APP degradation in APP-expressing cell models. To establish screening models, we tagged wild type APP695 (APP695), the predominant neuronal isoform^28^, with the small HiBiT tag (APP695-HiBiT) and expressed its in HEK293 cells. Cell lysates were analyzed using a luminescence-based assay of HiBiT-tagged protein^29^, identifying clone #3 and #9 with robust APP695-HiBiT expression (Extended Data Fig. 1a). Western blotting and ELISA confirmed intracellular expression of APP695 and extracellular release of Aβ42 in these clones (Extended Data Fig. 1b,c).

To assess CAPRIN1’s role in APP regulation, *CAPRIN1* DNA was overexpressed in HEK-APP695-HiBiT clones, leading to a dose-dependent decrease in intracellular APP and secreted Aβ42 (Fig. 1a,b). Co-transfection of HiBiT-APP695 and Flag-tagged CAPRIN1 in HEK cells, Flag immunoprecipitation (IP), followed by western blotting using HiBiT antibodies demonstrated the interaction between CAPRIN1 and APP (Fig. 1c). Conversely, CRISPR-Cas9 knockout of CAPRIN1 (sgCAPRIN1) increased both APP and Aβ42 levels (Fig. 1d,e), supporting CAPRIN1’s role in regulating APP degradation and Aβ42 production.

**Fig. 1.**
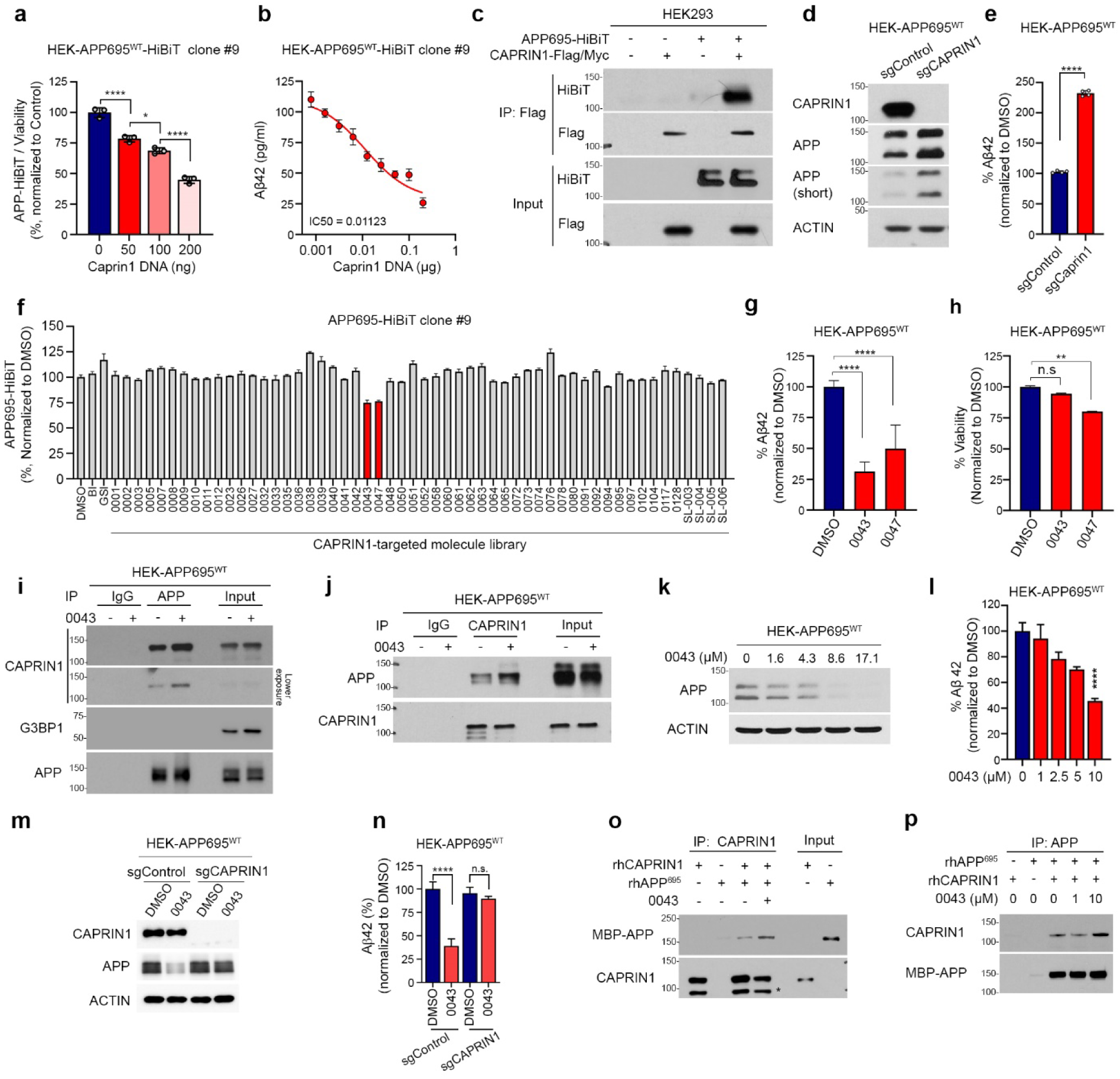
CAPRIN1 regulates APP levels and identifies compound 0043 as a small-molecule APP degrader. a, APP-HiBiT levels in HEK cells transfected with increasing doses of CAPRIN1 expression plasmid (mean ± s.d.; *n* = 3; one-way ANOVA with Tukey’s post hoc test). b, Dose–response curve for Aβ42 levels upon CAPRIN1 overexpression with calculated IC₅₀ (mean ± s.d.). c, Co-immunoprecipitation (IP) of CAPRIN1 and APP following co-transfection of APP695-HiBiT and CAPRIN1-Flag/Myc plasmids in HEK cells. d, Western blot analysis of APP and CAPRIN1 expression in sgCAPRIN1 HEK-APP695^WT^ cells. e, Aβ42 levels measured by ELISA in sgCAPRIN1 versus sgControl HEK-APP695^WT^ cells (mean ± s.d.). f, HiBiT signal in APP695-HiBiT clone #9 cells treated for 24 h with CAPRIN1-targeted small-molecule library (% of DMSO control). Compounds 0043 and 0047 are highlighted in red. g, h, HEK-APP695^WT^ cells treated with DMSO, 0043 or 0047 for 72 h. Aβ42 levels were measured by ELISA (g), and cell viability was assessed using CellTiter-Glo assay (h) (mean ± s.d.; *n* = 3; **p < 0.01, ****p < 0.0001; one-way ANOVA). i, j, IP showing enhanced APP–CAPRIN1 interaction in HEK-APP695^WT^ cells treated with 0043 (10 µM, 24 h); APP was immunoprecipitated in (i), CAPRIN1 in (j). k, Western blot of APP in HEK-APP695^WT^ cells treated with increasing concentrations of 0043 for 72 h. l, Aβ42 levels in culture media after 72 h treatment with 0043 (mean ± s.d.; *n* = 3). m, Western blot showing APP and CAPRIN1 expression in sgControl and sgCAPRIN1 HEK-APP695^WT^ cells treated with DMSO or 0043 (10 µM, 24 h). n, Aβ42 levels in the culture media of the same cells as in (m) after 72 h (mean ± s.d.; *n* = 3; ****p < 0.0001; one-way ANOVA with Tukey’s post hoc test). o, p, In vitro binding assays showing 0043 enhances the interaction between recombinant APP and CAPRIN1 (0043 = 10 µM).

Using these APP695-HiBiT clones, we screened our CAPRIN1-targeted compound library ^26^. Treatment with individual compounds for 24 hours reveal several compounds that reduced APP695-HiBiT levels, with compound 0043 and 0047 being more effective (Fig. 1f and Extended Data Fig. 1d). After 72-hour treatment, ELISA and cell viability assays indicated that compound 0043 was the most potent in reducing Aβ42 without inducing cytotoxicity (Fig. 1g,h and Extended Data Fig. 1e). These active compounds shared common pharmacophores: a bicyclic ring, a urea core, and a phenyl ring. The urea core was conserved in all the compounds. Regarding the bicyclic ring, the 6-5 ring system was most favorable for activity, with a benzothiazole ring as found in compound 0043, showing the higher potency (Extended Data Fig. 1f).

To assess target engagement, HEK-APP695 cells were treated with compound 0043; and APP and CAPRIN1 were isolated by IP using their respective antibodies. Western blot and ELISA showed that 0043 treatment enhanced the interaction between CAPRIN1 and APP (Fig. 1i,j) and reduced intracellular APP and extracellular Aβ42 levels in a dose dependent manner (Fig. 1k,l); whereas these effects were abolished in *CAPRIN1* KO cells (Fig. 1m,n). Notably, 0043 treatment had no effect on CAPRIN1 and its binding partner, Ras GTPase-activating protein-binding protein 1 (G3BP1) levels^30^, and qRT-PCR revealed no significant changes in APP mRNA (Extended Data Fig. 1g,h), supporting the post-translational mechanism of compound 0043.

To confirm direct protein-protein and compound interaction, His-tagged CAPRIN1 and MBP-tagged APP695 were purified and sedimented on glycerol gradients. SDS-PAGE showed a shift of peak fractions towards higher molecular weights when rhCAPRIN1 and rhAPP were mixed; and IP by CAPRIN1 antibodies, followed by western blots using MBP antibody, confirmed the direct interaction of CAPRIN1 and APP (Extended Data Fig. 1i-k). ThermoFluor assay showed thermal shifts for both proteins upon 0043 treatment, indicating compound binding of CAPRIN1 and APP (Extended Data Fig. 1l,m). Furthermore, compound 0043 enhanced reciprocal pull-down of CAPRIN1 and APP using specific antibodies (Fig. 1o,p). Together, these results identify compound 0043 as an APP degrader that enhances CAPRIN1-mediated APP degradation and reduces Aβ42 production in these cell models.

### The hit compound 0043 degrades intracellular APP in AD patient-derived iPSC-Ns

Human induced pluripotent stem cells (iPSCs) have accelerated disease modeling and targeted drug discovery^31^. The development of AD patient-derived iPSC neurons (iPSC-Ns) has provided relevant system to study this complex disease^32^. To assess compound activity in AD neurons, we differentiated iPSCs from sporadic AD (sAD), familial AD (fAD) and non-AD individuals into mutual neurons using doxycycline-inducible Neurogenic 2 (Ngn2) and Ascl1 (Msch1) expression ^33, 34^ (Extended Data Fig. 2a,b). Differentiated iPSC-Ns exhibited neuronal morphology, downregulation of stemness marker NANOG, and upregulation of neuronal marker MAP2 (Extended Data Fig. 2c,d). To examine CAPRIN1’s role in AD neurons, we immunoprecipitated APP from sAD-iPSC-Ns (AG27606-Ns) and confirmed its interaction with CAPRIN1 (Fig. 2a). No interactions were observed between CAPRIN1 and APP homologues: amyloid precursor-like proteins 1 and 2 (APLP1 and APLP2) (Extended Data Fig. 2e). *CAPRIN1* knockdown via its shRNAs increased intraneuronal APP and secreted Aβ42 (Fig. 2b,c), supporting the role of CAPRIN1 in APP degradation.

**Fig. 2.**
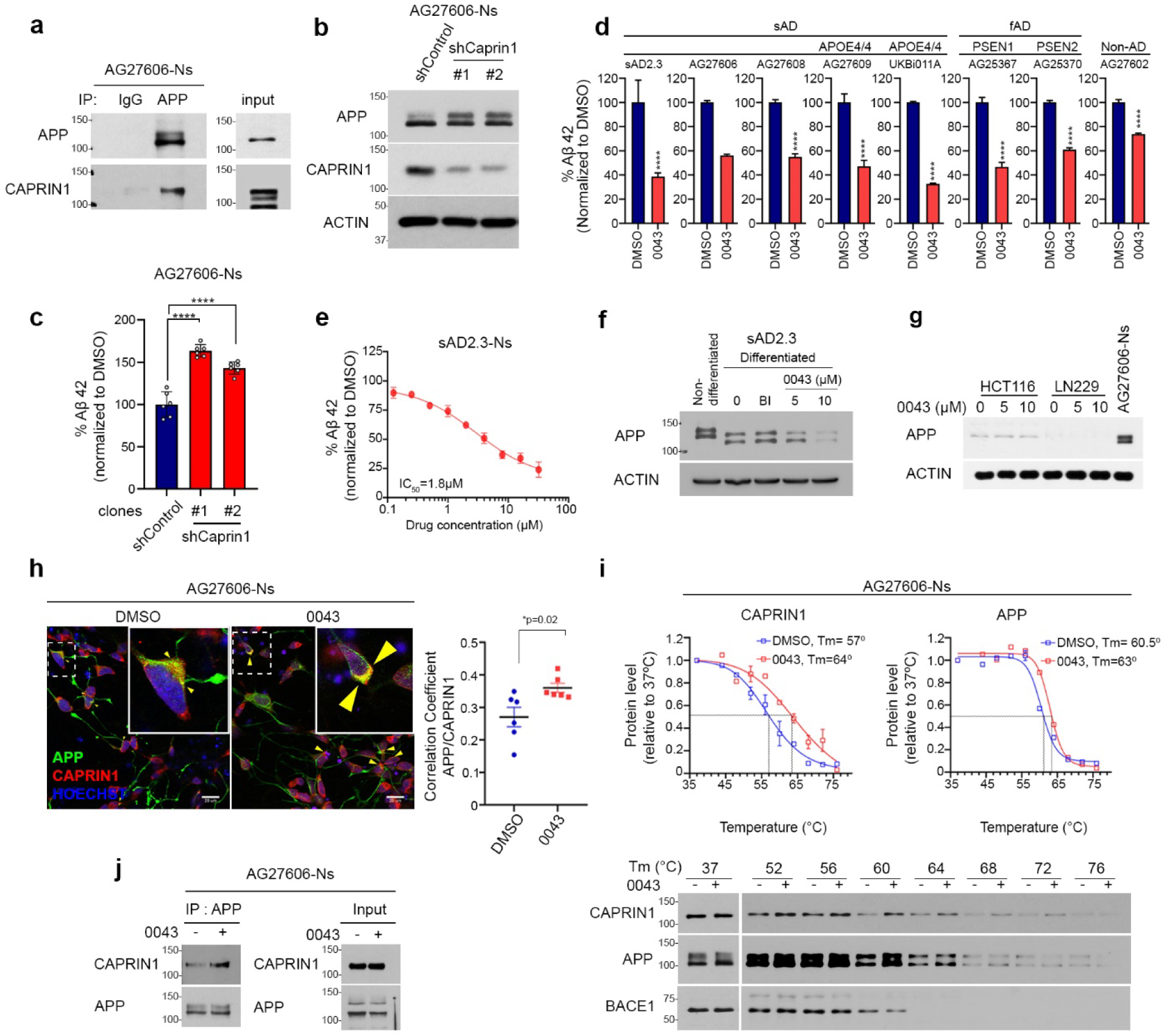
Compound 0043 promotes CAPRIN1-mediated APP degradation and reduces Aβ42 in AD iPSC-derived neurons. a, Endogenous interaction between APP and CAPRIN1 in AG27606 neurons detected by IP. b, Western blot of AG27606 neurons infected with shControl or shCAPRIN1 lentivirus for 72 h. c, Aβ42 levels measured by ELISA in culture media from the same cells as in (b) (mean ± s.d.). d, Aβ42 levels in culture media of multiple AD and non-AD iPSC-derived neurons treated with DMSO or 0043 (10 µM, 72 h) (mean ± s.d.; *n* = 4; ****p < 0.0001; unpaired two-tailed *t*-test). e, Dose–response curve of Aβ42 reduction in sAD2.3 neurons treated with 0043 for 5 days (IC₅₀ = 1.8 µM; mean ± s.d.; *n* = 3). f, Western blot showing APP levels in sAD2.3 neurons treated with 0043 (5 or 10 µM, 72 h); BACE1 inhibitor (BI) used as negative control. g, APP levels by western blot in HCT116 and LN229 cells treated with 0043 (10 µM, 72 h); AG27606 neurons used as neuronal control. h, Confocal images of AG27606 neurons treated with DMSO or 0043 (10 µM, 24 h), stained for APP (green), CAPRIN1 (red), and nuclei (Hoechst, blue). Arrowheads denote colocalization (scale bar, 20 µm). Quantification using Pearson’s correlation coefficient (*n* = 6; ***p < 0.001; unpaired two-tailed *t*-test). i, CETSA analysis of endogenous APP and CAPRIN1 in AG27606 neurons treated with 0043 (10 µM); thermal stability curves shown with BACE1 as control. j, IP of CAPRIN1 in AG27606 neurons treated with 0043 (10 µM, 24 h), followed by immunoblotting for APP.

We next evaluated compound activity and showed that both 0043 and 0047 reduced Aβ42 levels but 0047 was cytotoxic in sAD-iPSC-Ns (SAD2.3-Ns) (Extended Data Fig. 2f,g). Compound 0043 effectively lowered Aβ42 secretion across iPSC-Ns derived from sAD, fAD and non-AD iPSCs (Fig. 2d). Its activity was dose and time-dependent with IC_50_ values of 1.8-2.1 μM in SAD2.3-Ns and AG27606-Ns (Fig. 2e and Extended Data Fig. 2h,i). Western blotting confirmed that 0043 reduced intracellular APP, whereas the BACE1 inhibitor (BI, β-secretase inhibitor IV)^35^ did not in SAD2.3-Ns and AG27606-Ns (Fig. 2f and Extended Data Fig. 2j). Importantly, 0043 did not affect levels of other AD related proteins, including BACE1, ADAM10, PSEN1, nicastrin, ADP ribosylation factor-binding protein 1 (GGA1), Sorla, and neprilysin in SAD2.3-Ns, nor did it alter APP mRNA expression (Extended Data Fig. 2k,l). Compound 0043 degraded APP in iPSC-Ns but not in cancer cells that lack APP expression (Fig. 2g), indicating its neuronal-specific activity. To further assess the mechanism, fluorescence confocal microscopy showed increased co-localization of APP and CAPRIN1 after 0043 treatment in AG27606-Ns (Fig. 2h). Cellular thermal shift assay (CETSA)^36^ revealed that 0043 stabilized CAPRIN1 and APP but not BACE1 in both AG27606-Ns and HEK-APP695 cells (Fig. 2i and Extended Data Fig.2m). Finally, APP was purified from AG27606-Ns by IP after 0043 treatment; and western blotting revealed that the compound enhances CAPRIN1-APP interaction (Fig. 2j). Together, these findings demonstrate that compound 0043 binds CAPRIN1 and APP, enhancing their interaction and promoting CARPIN1-mediated degradation of APP in AD neurons.

### SAR studies of 0043 derivatives identify soluble and potent early lead compound 0152

Compound 0043 consists of a benzothiazole ring with morphine amide at R1 and a phenyl ring bearing a nitrile group at R2. It exhibited favorable drug-like properties in terms of molecular weight (MW), calculated partitional coefficient (cLogP), and polar surface area (PSA) but showed poor aqueous solubility by LC-MS analysis (Fig. 3a). To improve the solubility, we conducted initial structure-activity relationship (SAR) studies, introducing polar or flexible group and methyl substitutions of the urea linker to disrupt planarity (Extended Data Fig. 3a). A series of derivatives were designed, synthesized and screened in HEK-APP695-HiBiT cells and AG27606-Ns using luminescence, ELISA, and cell viability assay. Among the derivatives, compound 0152 emerged as a more potent and less cytotoxic analog, effectively reducing intracellular APP and extracellular Aβ42 without neuronal toxicity (Fig. 3b,c and Extended Data Fig. 3b). Compound 0152 was modified from compound 0043, replacing the nitrile with a methoxy group on the phenyl ring. It retained favorable MW, cLogP, and PSA), while significantly improved aqueous solubility (Fig. 3a). Central nervous system (CNS) multiparameter optimization (MPO) assessment^37^ suggested that compound 0152 had improved blood-brain barrier (BBB) permeability as compared to 0043. The MDR-MDCK assay showed that compound 0152 exhibited a lower efflux ratio than 0043, calculated as the ratio of the permeability coefficient in the apical to basolateral direction through the monolayers (Papp, A-B) (Fig. 3a), predicting improved oral absorption and brain penetration of compound 0152^38^.

**Fig. 3.**
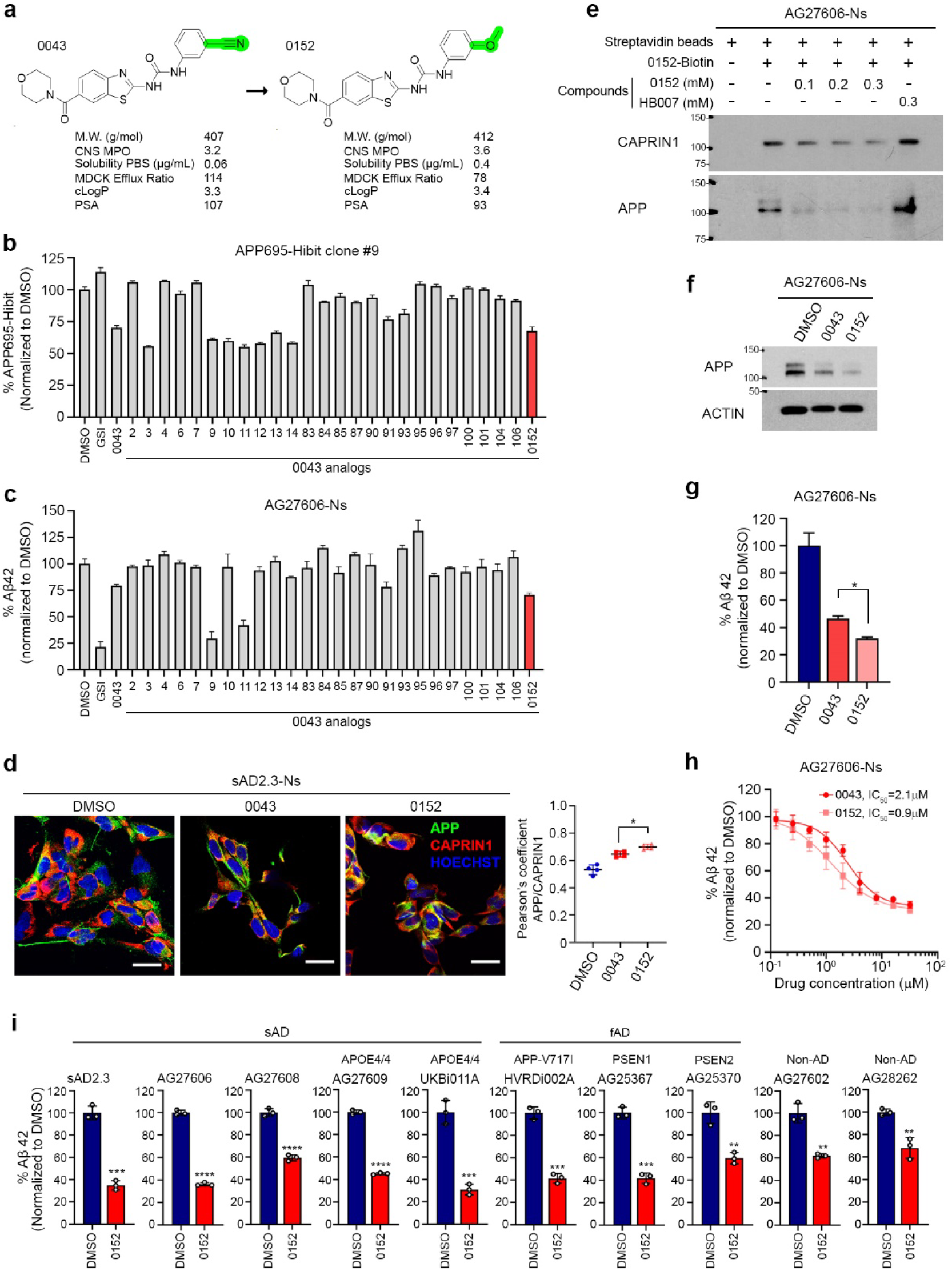
Compound 0152 is a more potent and soluble APP degrader than 0043. a, Structural comparison of 0043 and 0152, highlighting modifications and physicochemical properties (MW, CNS MPO, PBS solubility, MDCK efflux ratio, cLogP, PSA). b, APP695-HiBiT signal in HEK-APP695^WT^ cells treated with 0043 analogs (24 h), shown as % change from DMSO control; 0152 highlighted in red. c, Aβ42 levels in AG27606 neurons treated with 0043 analogs (72 h), shown as % change from DMSO control; 0152 highlighted in red. d, Confocal images and Pearson’s colocalization coefficient for APP and CAPRIN1 in sAD2.3 neurons treated with DMSO, 0043, or 0152 (10 µM, 24 h) (mean ± s.d.; *n* = 4; *p < 0.05). e, Streptavidin pull-down assay using biotinylated 0152 in AG27606 lysates. Competitive inhibition with unlabeled 0152 but not control compound HB007. f, Western blot showing APP reduction in AG27606 neurons after 5-day treatment with 0152 (10 µM). g, Aβ42 levels in AG27606 neurons treated with DMSO, 0043, or 0152 (10 µM, 72 h) (mean ± s.d.; *n* = 3; *p < 0.05, **p < 0.01, ****p < 0.0001; one-way ANOVA). h, Dose–response curves in AG27606 neurons after 5-day treatment with 0043 or 0152; 0152 showed improved potency (IC₅₀ = 0.9 µM vs. 2.1 µM; mean ± s.d.; *n* = 3). i, Aβ42 levels in culture media of iPSC-derived neurons from sAD, fAD (PSEN1/2, V717I), APOE4/4 carriers, and non-AD controls treated with 0152 (10 µM, 72 h) by ELISA (mean ± s.d., n = 3).

To examine compound binding to cellular CAPRIN1 and APP, we observed increased colocalization of APP and CAPRIN1 following treatment with either compound 0043 or 0152, with compound 0152 showing greater potency in AG27606-Ns and sAD2.3-Ns (Fig. 3d and Extended Data Fig. 3c). To assess direct target engagement, compound 0152 was chemically conjugated to biotin and biotin-conjugated 0152 (0152-biotin) was used in pull-down assays with AG27606-N lysates. Western blotting confirmed binding of 0152 to both CAPRIN1 and APP, which was competitively inhibited by excess free 0152 but not by the inactive analog HB007 (Fig. 3e). Functional assays showed that compound 0152 was more potent than 0043 in reducing both intracellular APP and extracellular Aβ42 (Fig. 3f–h and Extended Data Fig. 3d–f). Across a panel of iPSC-Ns derived from sAD, fAD, and non-AD individuals, compound 0152 consistently reduced both APP and Aβ42 levels (Fig. 3i and Extended Data Fig. 3g). Together, these findings identify compound 0152 as a promising APP molecular glue degrader with enhanced potency, solubility and BBB permeability.

### Compound 0152 functions as a molecular glue, forming a ternary complex with CAPRIN1 and APP

To quantify how compound 0152 enhances the CAPRIN1-APP interaction, we performed surface plasmon resonance (SPR) using recombinant proteins^39^. rhCAPRIN1 was immobilized on a CM5 sensor chip, and rhAPP695 was applied either before or after incubation with compound 0152. In the absence of the compound, SPR revealed a concentration-dependent interaction between APP695 and CAPRIN1. In the presence of equimolar 0152, the dissociation rate constant (K_off_) decreased 10-fold, while the association rate constant (K_on_) increased 3.5-fold, leading to a 36-fold reduction in the equilibrium dissociation constant (K_D_) (Fig. 4a-c). These kinetic changes indicate that compound 0152 stabilizes the CAPRIN1-APP interaction, consistent with a molecular glue mechanism.

**Fig. 4.**
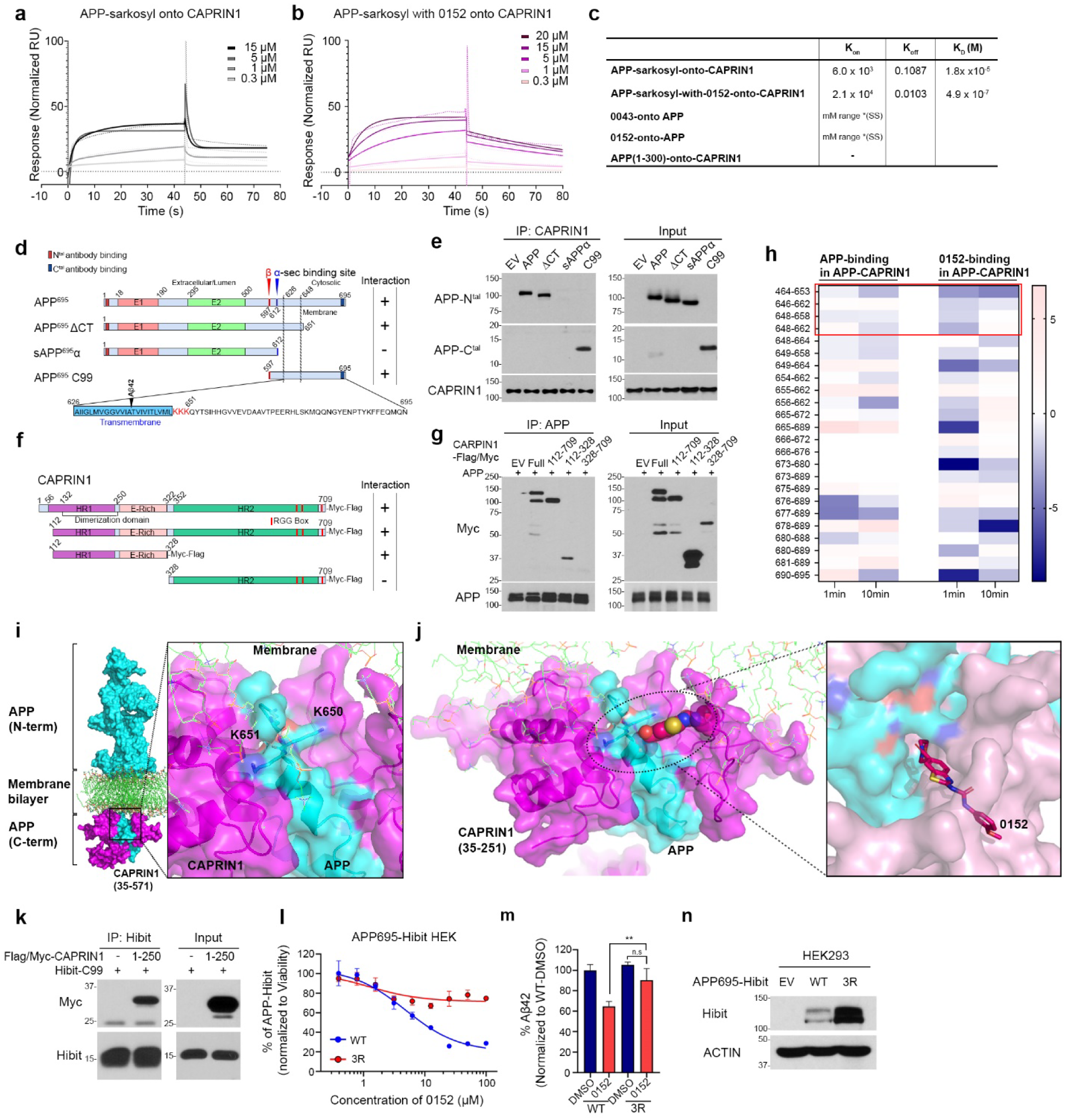
Compound 0152 stabilizes the APP–CAPRIN1 interface in vitro. a,b, SPR sensorgrams showing binding of sarkosyl-solubilized APP to immobilized CAPRIN1 in the absence (a) or presence (b) of 0152 (0.1 µM). Experimental data (dotted lines) and fits (solid lines) are shown for increasing APP concentrations. c, Kinetic parameters (K_D_: M; K_on_: M^-1^s ^-1^; K_off_: s^-1^) for APP–CAPRIN1 interactions with or without 0152; steady-state model used for 0043 and 0152 data. d, Schematic of APP fragments used for CAPRIN1 interaction mapping, indicating cleavage sites, transmembrane domain, and intracellular region. e, IP of APP fragments using CAPRIN1 antibody, detected by N-terminal or C-terminal APP antibodies. f, Diagram of CAPRIN1 truncation constructs (full-length, HR1, HR2, E-rich, RGG domains). g, APP IP with Flag/Myc-tagged CAPRIN1 truncations expressed in HEK293 cells. h, HDX-MS analysis of APP (residues 646–695) showing reduced deuterium uptake around the KKK motif (649–651) upon 0152 binding. Color scale indicates protection (blue) to deprotection (red). i, Structural model of the CAPRIN1–APP complex highlighting the APP KKK cluster (cyan) interacting with CAPRIN1 acidic surface (pink). j, Docking of 0152 at the CAPRIN1–APP interface. k, HiBiT pull-down assay showing reduced interaction between HiBiT-C99 and CAPRIN1(1– 250) in HEK293 cells. l, Dose–response HiBiT assay in APP695-HiBiT wild-type (WT) and 3R mutant cells after 72 h treatment (mean ± s.d.; n = 3). m, Aβ42 levels in culture media of WT and 3R cells treated with DMSO or 0152 (mean ± s.d.; n = 3; **p < 0.01; one-way ANOVA with Tukey’s post hoc test). n, Western blot showing reduced APP degradation in 3R mutant versus WT cells.

To map the interacting regions, we expressed domain constructs of APP and CAPRIN1 in HEK cells. Co-IP showed that the C-terminal region of APP interacts with CAPRIN1 (Fig. 4d,e), while the N-terminal domain of CAPRIN1 mediated APP binding (Fig. 4f,g). Hydrogen– deuterium exchange mass spectrometry (HDX-MS)^40, 41^ further defined the interaction interface. CAPRIN1 binding protected APP residues, including KKK motif (K649–K651), H658, Q679, and Y682 with the KKK motif showing the strongest protection (Fig. 4h and Extended Data Fig. 4a,b), CAPRIN1 regions 95–105, 160–170, and 295–430 showed reduced deuterium exchange upon APP binding, suggesting these regions as potential APP-interacting domains.

Molecular docking predicted a compact contact interface between the APP KKK motif and CAPRIN1 residues T103, L178, and S179 (Fig. 4i). Compound 0152 docked into this interface, forming hydrogen bonds with APP K650/K651 and CAPRIN1 Q90/D91 (Fig. 4i,j and Extended Data Fig. 4c,d). HDX-MS confirmed that compound 0152 enhanced protection of APP residue K650 and modest stabilization of surrounding residues (509–523), along with CAPRIN1 regions 95–105, 120–130, 245–255, and 292–314 (Extended Data Fig. 4e). Additional stabilization was attributed to helix–helix interactions between CAPRIN1 and the C-terminal domain of APP (Extended Data Fig. 4f). To validate the structural model, we co-expressed the C-terminal domain of APP (HiBiT-tagged C99) and N-terminus of FLAG-tagged CAPRIN1 (residues 1-250) in HEK cells. Flag IP followed by western blotting confirmed interaction between CAPRIN1 and APP, supporting the structural model of the ternary complex (Fig. 4k).

To assess the functional relevance of this interface, we generated an APP695-3R mutant, replacing the KKK motif (K649-K651) with arginines. HEK cell stably expressing 3R-APP695-HiBiT showed no response to compound 0152, as measured by HiBiT luminescence and Aβ42 ELISA (Fig. 4l,m). Western blotting of HEK cells transfected with equal amounts of wild-type (WT) or 3R-APP695-HiBiT DNA revealed increased expression of the 3R mutant as compared to wild-type APP, further supporting the critical role of the KKK motif in CAPRIN1-mediated APP degradation (Fig. 4n). Together, these results demonstrate that compound 0152 functions as a molecular glue by enhancing a ternary complex between CAPRIN1 and APP via the KKK motif, which is essential for APP degradation.

### APP degraders induce CAPRIN1-mediated lysosomal degradation of APP in iPSC-derived neurons

APP is synthesized at endoplasmic reticulum (ER)-associated ribosomes and trafficked through the secretory pathway to the plasma membrane, where it undergoes proteolytic cleavage to release Aβ peptides^42^. Once at the membrane, APP is internalized into early endosomes (EE), where it can either recycle via cycling endosomes or retrograde transport to the trans-Golgi network (TGN) or progress to late endosomes (LE) and lysosomes for degradation^43–46^ (Extended Data Fig. 5a). To determine how CAPRIN1 regulates APP degradation, we tracked its intracellular localization using fluorescent confocal microscopy with organelle-specific markers^47, 48^. In AG27606-Ns, APP colocalized with the EE antigen 1 (EEA1), LE marker Rab7, and the lysosomal associate membrane protein 1 (LAMP1) and treatment with compound 0043 significantly increased APP colocalization with all three markers (Extended Data Fig. 5b-d), suggesting that the compound enhanced APP trafficking through the endosomal-lysosomal pathway. CAPRIN1 also colocalized with Rab7 and LAMP1 and its colocalization was enhanced by compound 0043 (Extended Data Fig. 5e-g), indicating that the compound enhances the trafficking of CAPRIN1, like APP through the endosomal-lysosomal pathway.

Fluorescent confocal microscopy combined with coefficient analysis showed that both compounds 0043 and 0152 increased APP–LAMP1 colocalization, with 0152 showing greater potency across a panel of AD iPSC-Ns (Fig. 5a and Extended Data Fig. 5h,i). Importantly, CRISPR-mediated knockout of CAPRIN1 abolished this compound-induced colocalization, confirming CAPRIN1’s role in mediating APP trafficking to lysosomes (Fig. 5b and Extended Data Fig. 5j). Immunoprecipitation of CAPRIN1 following 0043 treatment revealed increased association with both APP and LAMP1 (Fig. 5c,d), supporting this trafficking mechanism. To further establish APP lysosomal degradation, we pretreated iPSC-Ns with the lysosome inhibitor Bafilomycin A1 (Baf A1) or the proteasome inhibitor MG132. MG132 had no effect, whereas Baf A1 abolished the APP-degrading activity of compounds 0043 and 0152 in AG27606-Ns, sAD2.3-Ns, and HEK-APP695 cells (Fig. 5e–h and Extended Data Fig. 5k,l). These findings indicate that compound-induced APP degradation is lysosome-dependent.

**Fig. 5.**
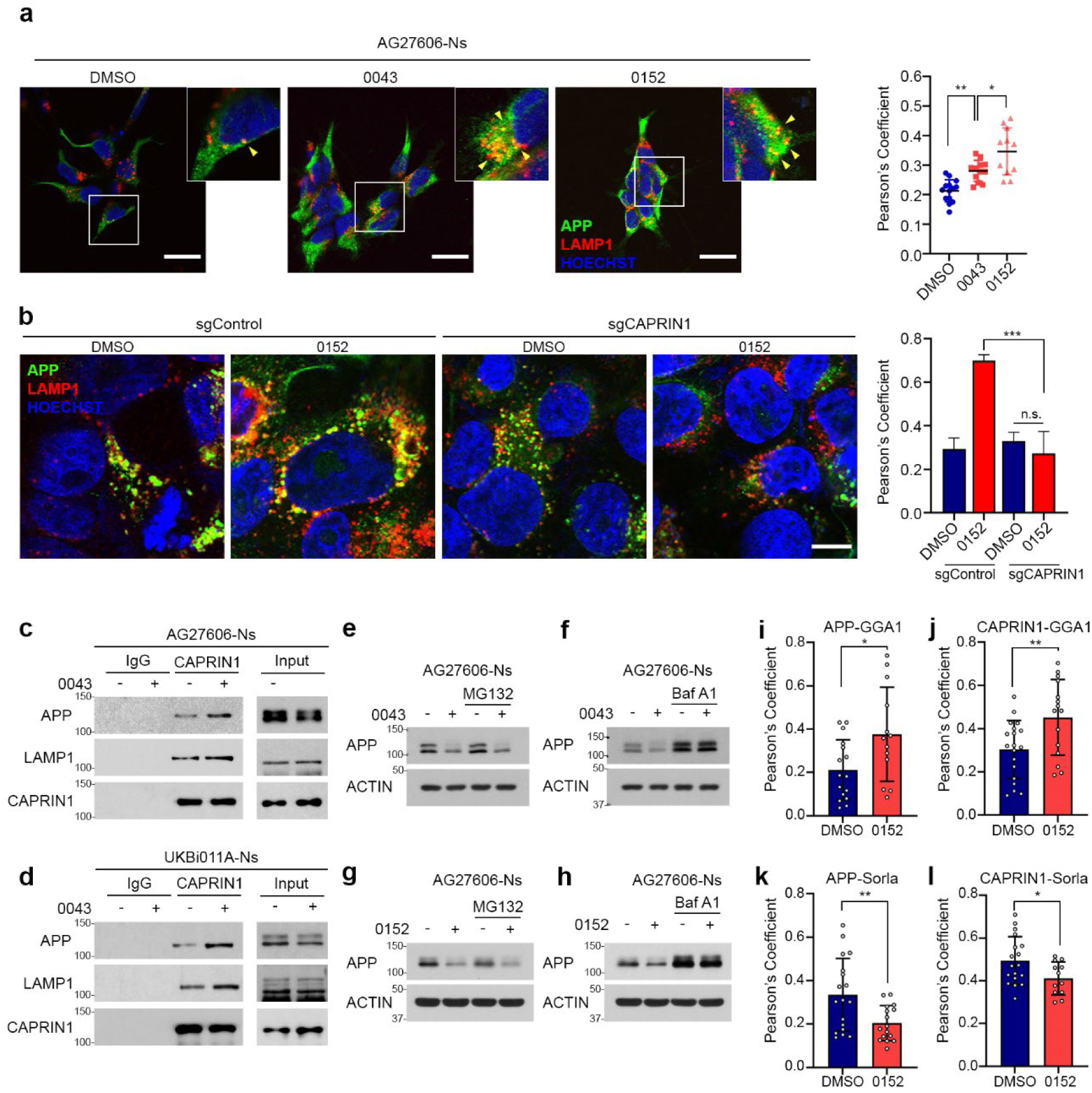
Small-molecule glue degraders direct APP to lysosomes via CAPRIN1. a, Confocal images of AG27606 neurons treated with DMSO, 0043, or 0152 (10 µM, 24 h), stained for APP (green), LAMP1 (red) and nuclei (Hoechst, blue). Yellow arrows indicate colocalization (scale bar, 20 µm). Right: quantification of APP–LAMP1 colocalization (mean ± s.d.; *p< 0.05; unpaired *t*-test). b, Confocal images of HEK-APP695^WT^ sgControl or sgCAPRIN1 cells treated with DMSO or 0152, stained for APP and LAMP1 (scale bar, 10 µm). Graph shows Pearson’s coefficient of APP–LAMP1 colocalization (mean ± s.d.; \*\**P* < 0.001). c, CAPRIN1 IP from AG27606 neurons treated with 0043, blotted for APP and LAMP1. d, APP and LAMP1 IP with CAPRIN1 in UKB011A neurons treated with 0043. e-h, Western blot of APP in AG27606 neurons pretreated with MG132 (5 µM, 30 min; e,g) or Bafilomycin A1 (20 nM, 30 min; f,h), followed by 0043 or 0152 (10 µM, 24 h). i,j, Pearson’s correlation of APP–GGA1 (i) and CAPRIN1–GGA1 (j) in AG27606 neurons treated with DMSO or 0152 (10 µM, 24 h, mean ± s.d.; *p < 0.05, **p < 0.01; unpaired *t*-test). k,l, Pearson’s correlation of APP–SORLA (k) and CAPRIN1–SORLA (l) after DMSO or 0152 treatment (10 µM, 24 h, mean ± s.d.; *p < 0.05, **p < 0.01).

We next examined whether compounds affect APP recycling via recycling endosome and TGN. Golgi-associated, gamma-adaptin ear-containing, ARF-binding protein 1 (GGA1) mediates sorting from endosomes to lysosomes^49, 50^ while sorting-related receptor with A-type repeats (SorLA) promotes recycling from endosomes back to the TGN^49, 51^. Fluorescent confocal microscopy revealed that 0152 treatment enhanced the colocalization of APP and CAPRIN1 with GGA1, which further confirms that the compound promotes CAPRIN1-mediated APP trafficking from endosomes to lysosomes (Fig. 5i,j and Extended Data Fig. 5m,n). In contrast, 0152 treatment reduced the colocalization of APP and CAPRIN1 with SorLA in AG27606-Ns (Fig. 5k,l and Extended Data Fig. 5o,p). These results suggest that compound 0152 redirects APP away from the recycling and retrograde pathway and toward CAPRIN1-mediated APP lysosomal degradation in AD neurons.

### CAPRIN1 and APP colocalization in neurons of AD brains and iPSC-brain organoids

To assess the role of CAPRIN1 in AD, we first examined its expression and localization in postmortem brain tissue. IHC of formalin-fixed, paraffin-embedded sections showed that CAPRIN1 was predominantly expressed in neurons of the cerebral cortex and hippocampus in AD and non-AD brains (Fig. 6a,b and Extended Data Fig. 6a,b). CAPRIN1 expression was absent in oligodendroglia and vascular endothelial cells, with weak positivity observed in a subset of astrocytes in both cerebral cortex and subcortical white matter (Fig. 6c and Extended Data Fig. 6c,d). CAPRIN1 was also detected in ependymal and choroid plexus epithelial cells (Extended Data Fig. 6d). As expected, APP was mainly neuronal and confocal immunofluorescence confirmed colocalization of CAPRIN1 and APP in neurons from AD brain sections (Fig. 6d and Extended Data Fig. 6e).

**Fig. 6.**
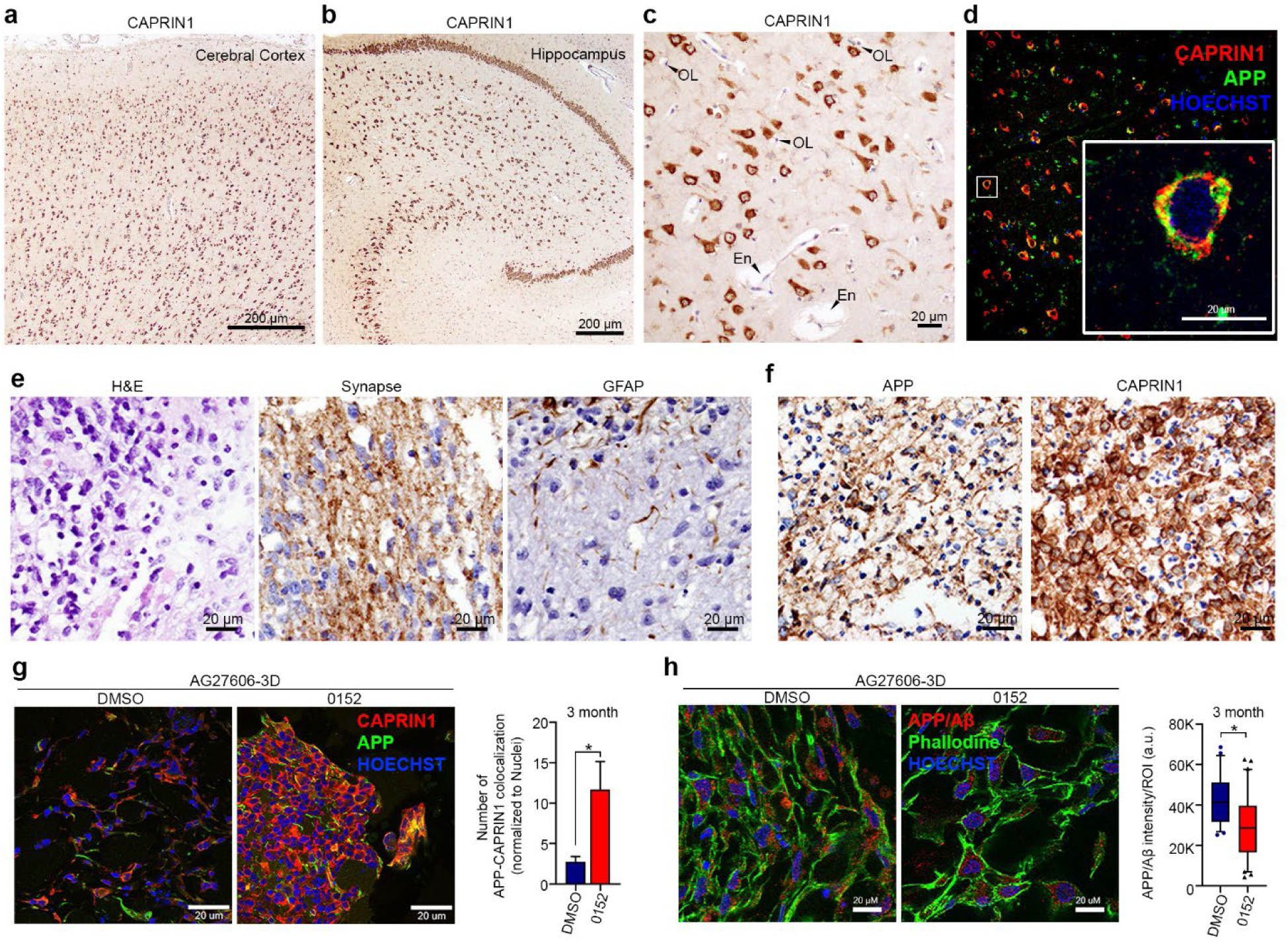
CAPRIN1 and APP co-localize in AD brain and organoids; 0152 reduces APP/Aβ levels in 3D cortical models. a,b, IHC of CAPRIN1 in postmortem AD brain cortex (a) and hippocampus (b), showing cytoplasmic localization (scale bars, 200 µm). c, CAPRIN1 in oligodendrocytes (OL) and endothelial cells (En) in AD cortex (scale bar, 20 µm). d, Confocal immunofluorescence of APP (green) and CAPRIN1 (red) with Hoechst (blue) in human AD brain (scale bar, 20 µm). e, H&E and IHC for synaptophysin and GFAP in AG27606 cortical organoids (scale bar, 20 µm). f, IHC of APP and CAPRIN1 in 12-week-old organoids (scale bar, 20 µm). g, Confocal images of 3-month AG27606 organoids treated with DMSO or 0152 for 3 months; stained for APP (green), CAPRIN1 (red), Hoechst (blue) (scale bar, 20 µm). Right: quantification of APP–CAPRIN1 colocalization (mean ± s.d.; *p < 0.05). h, Confocal images of organoids stained for APP/Aβ (red), phalloidin (green) and Hoechst (blue) after DMSO or 0152; quantification of APP/Aβ intensity (mean ± s.d.; *p < 0.05).

To model AD brain tissue and assess compound activity, we generated 3D brain organoids from patient-derived iPSCs (AG27606 and UKBi011A) using established protocols^52, 53^. Ngn2/Ascl1-iPSCs were cultured to form embryoid bodies (EBs) for five days, induced with doxycycline for five days, embedded in Matrigel and matured for 12 weeks (Extended Data Fig. 6f). Hematoxylin and eosin (H&E) and IHC confirmed neuronal and astrocytic differentiation, marked by the expression of synaptophysin and glial fibrillary acidic protein (GFAP), respectively (Fig. 6e). IHC further showed the expression of CAPRIN1 and APP in neurons in the organoids (Fig. 6f). After the 12-week maturation, organoids were treated with compound 0152 for an additional three months. Confocal microscopy and colocalization coefficient analysis of frozen sections showed that 0152 increased CAPRIN1–APP colocalization in organoids (Fig. 6g). Furthermore, staining with an antibody recognizing both APP and Aβ42 revealed that 0152 treatment reduced APP/Aβ42 levels in organoids derived from both AG27606 and UKBi011A iPSCs (Fig. 6h and Extended Data Fig. 6g). These findings demonstrate that CAPRIN1 and APP are co-expressed and colocalized in neurons of both AD brains and patient-derived organoids. Importantly, compound 0152 enhances this protein-protein interaction and reduces APP/Aβ42 accumulation in 3D AD organoids.

### Compound 0152 reduces Aβ and amyloid plaques in AD mouse brains

To evaluate in vivo activity, we first assessed the pharmacokinetics (PK) of compounds 0043 and 0152. Mice were dosed by oral gavage or intravenous injection, and plasma samples were collected over time for LC-MS analysis. PK profiling showed that compound 0152 achieved significantly greater drug exposure, with an oral bioavailability (F%) reaching up to 30% compared to only 5% for 0043 (Fig. 7a,b), indicating improved drug-like properties of 0152 compared to 0043. In evaluation of BBB permeability, oral administration of 0152 demonstrated its BBB permeability as evidenced by a favorable brain-to-plasma (B/P) partition co-efficient (Kp)^54^.

**Fig. 7.**
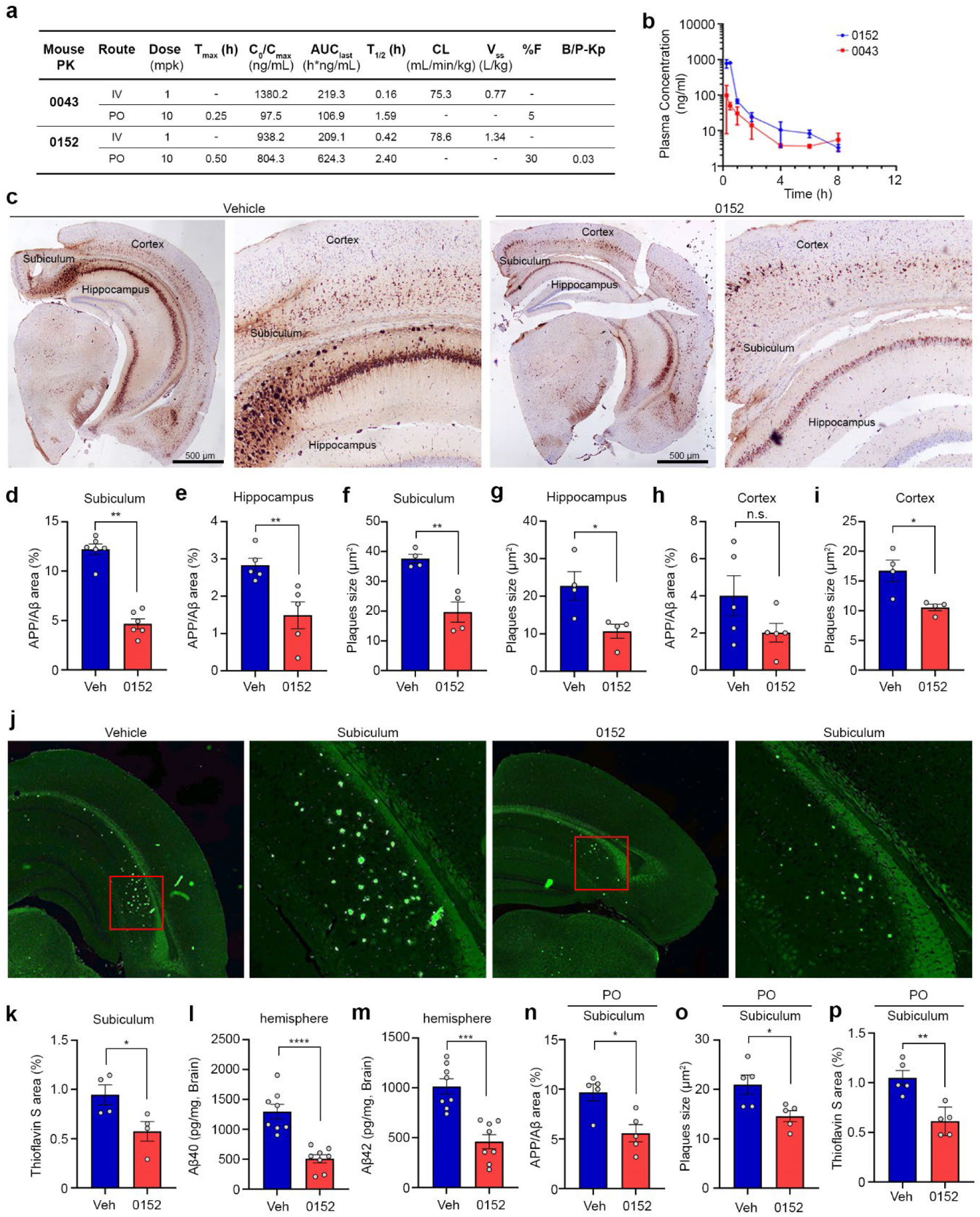
Compound 0152 reduces amyloid burden in 5xFAD mice. a, Pharmacokinetic parameters of 0043 and 0152 after intravenous (IV) and oral (PO) administration, showing improved bioavailability (%F), half-life (T₁/₂), and clearance (CL) for 0152. b, Plasma concentration–time profiles following single PO dose (100 mg/kg) in mice (*n* = 3 per group). c, IHC images of APP/Aβ accumulation in cortex, hippocampus, and subiculum after 24-day treatment with vehicle or 0152 (100 mg/kg, IP); representative whole-brain and subiculum sections shown (scale bars, 500 µm). d,e, Quantification of APP/Aβ-positive area (%) in subiculum (d) and hippocampus (e) of 5xFAD mice treated with 0152 (100 mg/kg, IP). f,g, Quantification of individual plaque size in subiculum (f) and hippocampus (g). h,i, Quantification of APP/Aβ area (h) and plaque size (i) in cortex. j,k, Thioflavin S staining of brain sections (j) and quantification in subiculum (k) of vehicle- and 0152-treated mice (*p < 0.05; unpaired *t*-test). l,m, Quantification of Aβ40 (l) and Aβ42 (m) in hemispheric brain lysates from 5xFAD mice treated with vehicle or 0152 (100 mg/kg, IP), measured by ELISA. mean ± s.e.m.; ***P < 0.001, ****P < 0.0001; unpaired t-test. n-p, Subicular amyloid pathology in 5xFAD mice (vehicle vs. 0152, 80 mg/kg, PO, 30 days): (n) APP/Aβ area, (o) plaque size, (p) ThioS area. Mean ± s.e.m.; *P < 0.05; unpaired t-test.

To assess effects on amyloid burden, we used nine-week-old 5xFAD mice, which start to rapidly accumulate Aβ and develop amyloid plaques from 8 weeks due to expression of mutant human APP and PS1 genes^55^. 5xFAD mice were subjected to three different treatment protocols. In the first protocol, 9-week-old 5xFAD mice received daily intraperitoneal injections of 100 mg/kg 0152 for 24 days. IHC of brain sections using anti-APP antibody revealed significant reductions in APP/Aβ-positive area and amyloid plaque size in the hippocampus and subiculum (Fig. 7c–g). The cortex, where amyloid deposition occurs later^55^, showed minimal changes in plaque burden (Fig. 7h,i). Thioflavin S staining confirmed reduced fibrillar Aβ plaques in the subiculum (Fig. 7j,k). ELISA of brain lysates corroborated the IHC results, showing marked decreases in both Aβ40 and Aβ42 levels in the hemispheres (Fig. 7l,m). Throughout the treatment period, mice tolerated treatment well, with no observed signs of lethargy, ataxia, paralysis, seizures, or weight loss (Extended Data Fig. 7a).

In a second experiment, 9-week-old 5xFAD mice were injected intraperitoneally with 80 mg/kg of 0152 twice per day for 30 days, which was well tolerated with no weight loss (Extended Data Fig. 7b). IHC analysis confirmed that 0152 treatment significantly reduced APP/Aβ-positive areas and amyloid plaque sizes in the subiculum and hippocampus, as well as amyloid plaque sizes in the cortex (Extended Data Fig. 7c–i). Thioflavin S staining further demonstrated that 0152 treatment drastically reduced fibrillar Aβ plaques in the subiculum (Extended Data Fig. 7j,k). In a third protocol, 9-week-old 5xFAD mice were treated via oral gavage with 80 mg/kg of 0152 twice daily for 30 days. IHC and Thioflavin S staining demonstrated a significant reduction of APP/Aβ-positive areas and plaque sizes in the subiculum of 0152-treated mice compared to controls with no observed weight loss (Fig. 7n–p and Extended Data Fig. 7m–o). Collectively, these findings provide strong in vivo evidence that compound 0152 reduces Aβ accumulation and amyloid plaque formation in the 5xFAD AD mouse model.

## DISCUSSION

Over the past decades, therapeutic strategies for AD have primarily focused on inhibiting BACE1 and γ-secretase to reduce Aβ peptide production^56^. However, clinical trials of BACE1 and γ-secretase inhibitors have encountered significant setbacks due to limited efficacy and off-target effects, as these enzymes are involved in physiological processes beyond Aβ generation^57^. More recently, mAbs targeting Aβ have been developed to facilitate amyloid clearance. Clinical trials of Aβ-directed mAbs, including aducanumab, lecanemab, and donanemab, have demonstrated their ability to reduce amyloid burden and modestly slow cognitive decline in AD patients at early stage. Nonetheless, these treatments are associated with substantial risks such as cerebral hemorrhage and amyloid-related imaging abnormalities (ARIA), highlighting the urgent need for safer and more effective therapeutic strategies^58^.

Parallel to efforts in the development of Aβ-targeted therapies, accumulating evidence has identified neuronal overexpression of APP as a central driver of Aβ pathology, positioning APP as a promising therapeutic target in AD, ADRD, and brain aging^11–15^. Antisense oligonucleotides (ASOs) have recently emerged as a strategy to modulate *APP* mRNA, resulting in decreased APP expression and reduced Aβ production in iPSC-derived neurons^59^. The clinical success of ASO therapies in spinal muscular atrophy provides a strong translational precedent^60^, and early-phase trials of intrathecal tau-targeting ASOs in patients with mild AD have shown encouraging results^61^. Despite these advances, ASO-based strategies face a significant limitation in AD; poor penetration of the BBB, which restricts delivery to deep brain tissues^62, 63^. In contrast, we demonstrate that APP-targeted molecular glue degraders reduce intraneuronal expression of APP and extracellular release of Aβ, and upon oral administration, the lead compound 0152 can cross the BBB and significantly reduce Aβ production and amyloid burden in the brains of 5xFAD mice.

Molecular glue degraders harness endogenous degradation pathways, primarily the ubiquitin–proteasome system for cytosolic proteins and the endo-lysosomal system for membrane-associated proteins, to selectively reduce or eliminate pathogenic proteins^21–23^. Advances in computational ligand-based drug design have enabled the development of homologous compound series that interface with a common E3 ligase and multiple target proteins, effectively expanding the "degradable proteome" or "degrome”^64, 65^. Previously, we reported HB007, a molecular glue degrader that targets CAPRIN1 and FBXO42, which mediates the recruitment of the FBXO42 substrate SUMO1 to the FBXO42-CUL1 E3 ligase and proteasomal degradation in cancer^26^. In the present study, we discovered a previously unrecognized role of CAPRIN1 in neurons: facilitating lysosomal degradation of APP through direct interaction. Screening our CAPRIN1-targeted compound library, we identified hit compound 0043 and optimized it into lead compound 0152. These compounds bind at the CAPRIN1–APP interface, stabilize the ternary complex, and promote CAPRIN1-mediated lysosomal degradation of membrane-bound APP in AD neurons. Notably, the distinct cellular activities of our SUMO1 and APP degraders likely reflect the cell-type-specific expression of their respective targets, in which APP is not expressed in cancer cells, while FBXO42–CUL1 complex is largely absent in neurons^26^.

CAPRIN1 is a neuronal RNA-binding protein involved in synaptogenesis, brain development, and neurodegeneration^66^. It forms a stable complex with G3BP1 that is essential for synapse formation^67^, and genetic deletion of either *Caprin1* or *G3bp1* in mice results in perinatal lethality due to impaired synaptic and respiratory function^68, 69^. CAPRIN1 also regulates RNA stress granules and has been implicated in neurodegenerative diseases, including AD^70^. The data uncover a novel functional role for CAPRIN1 in regulation of APP homeostasis in neurons. APP is synthesized in the endoplasmic reticulum and trafficked to the plasma membrane, where it undergoes sequential cleavage to release Aβ peptides^42^. Following endocytosis, membrane-bound APP is routed to the endo-lysosomal system for degradation^71^. We show there that CAPRIN1 is predominantly expressed in cortical and hippocampal neurons of the human brain. CAPRIN1 interacts specifically with membrane-bound APP via its cytoplasmic KKK motif and facilitates APP trafficking from endosomes to lysosomes for degradation. This interaction appears to be selective, as no binding was observed between CAPRIN1 and the APP homologues APLP1 or APLP2. Our compounds bind CAPRIN1 and APP, stabilize their interaction and enhance the endo-lysosomal degradation of membrane-bound APP without affecting cytosolic CAPRIN1, consistent with the multifunctional role of CAPRIN1 in neurons.

This study provides proof-of-concept for a CAPRIN1-dependent APP degradation and establishes CAPRIN1-targeted molecular glues as a novel therapeutic strategy for AD. Using our TPD-based drug discovery platform^26^, we developed cell-based screening assays that identified compound 0043 as the first APP degrader. Medicinal chemistry optimization yielded compound 0152, which exhibited improved potency, solubility, and BBB permeability, supporting its suitability for in vivo application. These compounds promoted CAPRIN1-mediated lysosomal degradation of wild-type and mutant APP in neurons derived from AD patient iPSCs. In vivo, treatment with compound 0152 significantly reduced Aβ accumulation and amyloid plaque formation in the brains of 5xFAD mice. Ongoing efforts are focused on optimizing the pharmacokinetic and pharmacodynamic profiles of these compounds using both ligand-based and structure-based drug design approaches to further enhance their potency and drug-like properties for clinical development. Collectively, our findings uncover a previously unrecognized CAPRIN1-mediated lysosomal degradation pathway for APP and introduce a first-in-class series of APP-targeted molecular glue degraders. This strategy represents a promising therapeutic avenue for AD, with the potential to overcome limitations associated with existing Aβ-directed therapies.

## Acknowledgements

This work was supported in part by the National Institute on Aging (R41 AG090241, C.H.) and the National Institute of Neurological Disorders and Stroke (R01 NS126358, C.H.). C.H. was supported in part by the endowment fund of the Bicentennial Chair at the Indiana University School of Medicine. The authors acknowledge the assistance of the Chemical Genomics Core and the Indiana Center for Biological Microscopy of the Indiana University School of Medicine. The authors also acknowledge the Bio-Instrumentation Core (BIC) at Notre Dame University for allowing them to perform the SPR experiments using the Biacore T200 system, and the mass spectrometry facility at Rutgers University for conducting the HDX-MS experiments.

## Author contributions

C.H., A.C.B. and A.S. initiated this APP drug discovery project. C.H., A.C.B. and S.H.J. designed the experiments and provided supervision. S.H.J. carried out all the drug screenings, molecular, biochemical, iPSC cellular experiments, and animal studies. R.R., N.J., R.K.R., Y.T. R.K.H. expressed and purified recombinant proteins. R.R. designed and performed structural studies. N.J. designed and carried out recombinant protein interaction. B.L., X.W., C.G., L.Z., and H.Y.L. designed, synthesized, and analyzed compounds. R.K.W. assisted biological work. S.H.J., A.C.B. and C.H. analyzed and organized all the data and co-wrote the manuscript. All authors reviewed and contributed to editing the manuscript.

## Online Methods

### Stable APP695 HEK293 cells

HEK293 cells were purchased from the American Type Culture Collection (ATCC). Lentiviral plasmids (pLV-Hygro-CMV-hAPP, pLV-Hygro-CMV-hAPP-Hibit, and pLV-Hygro-CMV-hAPP -3K to 3R-Hibit) for APP695^WT^ HEK293 and APP695-Hibit HEK293 were generated in the 3^rd^ generation plasmid. HEK293T cells were transfected with targeting plasmid and 3^rd^ generation packaging plasmid (pMD2.G; addgene#12259, pMDLg/pRRE; Addgene#12251, and pRSV-Rev; Addgene #12253) together using Lipofectamine 2000. HEK293 cells were infected with virus particles and selected with hygromycin B (200 µg/ml, Invitrogen). For single clone selection, the cells were reseeded in 96-well plates as single cells. Once expanded, clones were analyzed by ELISA for amyloid beta 42 (Aβ42), western blot for APP expression, and Hibit-lytic assay (Promega). For sgCAPRIN1 HEK293 cell lines, the guide RNA sequences of CAPRIN1 were cloned into LentiCRISPRv2 plasmid (Addgene #52961) according to protocol ^72^ and the sequences for CAPRIN1-CRISPR (in Supplementary Table 1). HEK293 cells were infected, selected with puromycin (1 µg/ml) for 2 days, single-cell selected, and followed by western blot for CAPRIN1 knock-out verification.

### Human iPSCs and Differentiation of iPSCs into Neurons

The human iPSC SAD2.3 (UCSD234i-SAD2-3) and HVRDi002A were purchased from Wicell, UKBi011A from European Bank for induced pluripotent stem cells, Sigma, AG25367, AG25370, AG27606, AG27608, AG27609, AG28049, AG28262, and AG27602 from Coriell. All iPSCs were maintained and expanded on Matrigel (Growth factor reduced Matrigel matrix, Corning) in mTeSR1 complete medium (mTeSR1 medium + 5X supplement, Stem Cell Technologies). For iPSC-Ns transgene delivery, each iPSC line was transduced with lentiviral particles for pLVX-UbC-rtTA-Ngn2:2A:Ascl1 (Addgene #127289) ^73, 74^, followed by selection in the presence of puromycin (0.5 mg/ml, Gibco). To initiate conversion, iPSCs were differentiated into neurons using the established Ascl1/Mash1^33^ and Neurogenin 2 (Ngn2)^75^ protocols. The transgene-delivered iPSCs were dissociated with StemPro™ Accutase™ (Gibco) and reseeded in Neuro culture media (Neurobasal media (Gibco), 2% B27 supplement (Gibco), 1X Glutamax (Gibco)) with doxycycline (2 µg/ml, Sigma-Aldrich) for 5 days and without doxycycline for 3 days^76^ and differentiated iNS exhibited differentiated neuronal morphology.

### iPSC-derived brain organoids

For generating brain 3D organoids, the Lancaster’s protocol ^52^ and Paşca’s protocol ^53^ were used with minor modifications. In brief, 9,000 cells/ 100 µL of each well were seeded in Ultra-low attachment 96 well U-bottom plates (Sbio) supplemented with 10 µM of ROCK inhibitor. The next day, 100 µl of mTeSR1 was added and the cells were cultured as embryonic bodies (EB) for 5 days at 37°C in 5% CO_2_ incubator. When the size of EB reached 400-600 µm, EBs were transferred into ultra-low binding 24 well (Corning) with 200 µl of neural differentiation medium (Neurobasal medium (Gibco), 2% B27, 1X Glutamax) with 2 µg/ml of Doxycycline and incubated for 5 days at 37°C in 5% CO_2_ incubator until the size of EB was 700-800 µm. During this step, one EB was in one well and the culture medium was changed every day with fresh neural differentiation medium with Doxycycline. At day 10, the EB was embedded into 15 µl of 100% Matrigel (Corning) and the embedded EBs were cultured in neural differentiation medium without Doxycycline in 6 well ultra-low binding plates (Corning). In this step, each well of 6 well plates had 16 EBs, and plates were incubated in stationary methods for 5 days at 37°C in 5% CO_2_ incubator to expand neuroepithelial buds, followed by placing the plates on the orbital shaker (Benchmark Scientific) at 75 rpm at 37°C in 5% CO_2_ incubator. The fresh medium was changed every 3 days.

### Animal experiments

Female 5xFAD transgenic mice (9–10 weeks old) were used to evaluate the efficacy of 0152 in reducing amyloid pathology. Mice were housed under Indiana University standard laboratory conditions. For intraperitoneal (IP) administration, mice received either vehicle or 0152 (indicated concentrations) in a formulation consisting of 20% DMSO, 20% Cremophor EL, and 60% PBS. For oral (PO) administration, mice received either vehicle or 0152 (indicated concentrations) in a formulation consisting of 10% NMP (N-methyl-2-pyrrolidone), 10% PG (Propylene Glycol), 5% Solutol HS-15, and 75% Captisol (10% w/v in RO water). At the end of the treatment period, mice were sacrificed, and their brains were collected for histological analysis. Brain tissues were fixed in 10% Formalin, embedded in paraffin, and sectioned. Thioflavin S staining was performed to visualize β-sheet-rich amyloid plaques, and immunohistochemistry was conducted using the 6E10 monoclonal antibody to detect Aβ plagues. Body weights were recorded daily throughout the study.

### PK studies

IV and PO PK in rodents were processed by Sai Life Sciences Pvt. Ltd using 8-to 10-week-old C57BL/6 male mice following a single intravenous administration at 1 mg/kg and oral administration at 10 mg/kg. A total eighteen mice (n=9/group) were divided into two groups as Group 1 and Group 2 with sparse sampling design (3 mice/time points). Animals in Group 1 were administered intravenously as slow bolus injections through tail vein at 1 mg/kg dose and animals in Group 2 were administered through oral route at 10 mg/kg. For 0043, the formulation vehicle used for IV group was 5% NMP, 10% PG, 5% Solutol HS-15 and 80% Captisol (10% w/v in RO water) and the formulation used for PO group was 10% NMP, 10% PG, 10% TPGS and 70% Captisol (10% w/v in RO water). For 0152, the formulation vehicle used for IV group was 5% DMSO, 20% Solutol HS-15 and 75% PBS and the formulation used for PO group was 5% NMP, 10% PG, 5% Solutol HS-15 and 80% Captisol (10% w/v in RO water). Blood samples (approximately 60 µL) were collected under light isoflurane anesthesia (Surgivet®) from retro orbital plexus from a set of three mice at 0.08, 0.25, 0.5, 1, 2, 4, 6, 8 and 24 hours for IV and PD, 0.5, 1, 2, 4, 6, 8 and 24 hours for PO. Immediately after blood collection, plasma was harvested by centrifugation at 10,000 rpm, 10 min at 40 °C, and samples were stored at -70±10°C until bioanalysis. All samples were processed for analysis by protein precipitation method and analyzed with fit-for-purpose LC-MS/MS method (LLOQ = 1.01 ng/mL). The plasma pharmacokinetic parameters were estimated using non-compartmental analysis tool of Phoenix® WinNonlin software (Ver 8.3).

### Brain/Plasma(B/P) Ratio test

Brain/Plasma Ratio test was processed by Sai Life Sciences Pvt. Ltd. Briefly, twelve male C57BL/6 mice were administered orally with solution formulation of 0152 at 10 mg/kg dose. Blood samples (approximately 60 µL) were collected under light isoflurane anesthesia from a set of three mice at 0.25, 0.5, 1 and 2 h. The blood samples were collected at each time point in labelled micro centrifuge tube containing K_2_EDTA as anticoagulant. Plasma was harvested by centrifugation of blood and stored at -70±10°C until analysis. Immediately after collecting blood, animals were anesthetized, perfused using 10 mL phosphate buffer saline and brain samples were collected from each mouse at respective time points. Brain samples were homogenized using ice-cold phosphate buffer saline (pH 7.4) and homogenates were stored below - 70±10°C until analysis. Total homogenate volume was three times the brain weight. The plasma and brain concentration-time data of 0152 were used for calculation. Plasma and brain samples were quantified by fit-for-purpose LC-MS/MS method.

### MDR1-MDCKII Assay

MDR1-MDCKII assay was processed by Sai Life Sciences Pvt. Ltd. Briefly, MDR1-MDCKII cells were seeded onto the polycarbonate membranes in the 24-well insert system at 0.12 x 10^6^ cells/well and cultured for 8 days until confluence before being used for the transport studies. Test compounds were diluted with the transport buffer (HPSS with 10 mM HEPES, pH 7.4) from DMSO stock solution to a concentration of 2 µM (DMSO < 1%) and applied to the apical or basolateral side of the cell monolayer. The plate was incubated for 2.0 h in CO_2_ incubator at 37±1 °C, with 5% CO_2_ at saturated humidity without shaking. Permeation of the test compounds from A to B or B to A direction was determined in duplicate. Loperamide (P-gp efflux substrate) was tested at 2 µM bidirectionally, Verapamil was used as P-gp inhibitor. While Atenolol (low permeability marker) and Propranolol (high permeability marker) were tested at 2.00 µM in A to B direction in duplicate. Test and reference compounds were quantified by LC-MS/MS analysis based on the peak area ratio of analyte/internal standard (IS). The efflux ratio of each compound was calculated. After the transport assay, Lucifer yellow fluorescence rejection assay was performed to confirm the integrity of the cell monolayer.

### Kinetic Solubility Assay

10 µL of 10 mM DMSO stock solution of test and control compounds was added to 990 µL phosphate buffer with pH 7.4. The solubility samples were vortexed for at least 2 minutes, then shaken on a shaker at room temperature for 24 hours. Samples were centrifuged, and the supernatant was transferred into a filter plate. Then, the filtrates were quantified by LC-MS system to get solubility results.

### Cellular Thermal Shift Assay (CETSA)

For CETSA, as described previously ^36^, iPSC-Ns or APP695^WT^-HEK were treated in the presence or absence of 0043 (100 µM) for 1 hour, trypsinized, washed, and resuspended in 100 µl of PBS with protease inhibitors (600,000 cells/tube). Cells were incubated at their designated temperatures for 3 minutes, followed by 25°C for 3 minutes. Then, NP40 (Final 1%) was added to lyse the cells, followed by immediate snap freezing with liquid nitrogen and thawing using a thermal cycler at 25°C. This freezing and thawing cycle was repeated twice. The lysates were spun at 13,500 rpm for 20 minutes at 4°C and the supernatant was harvested for western blotting analysis.

### Thermofluor Assay

For the Thermofluor assay, Huynh’s protocol ^77^ was used with minor modifications. In brief, all proteins were used at a final concentration of 5 µM for this assay, and GloMelt™ Thermal Shift Protein Stability Kit (Biotium) was used at a final concentration of 0.5X. All experiments were carried out with QuantStudio 6 Flex (Applied Biosystems). SYBR was used as a reporter and ROX was used as a passive reference according to the manufacturer’s directions. Melting curve data was exported, followed by analysis with Boltzmann’s sigmoidal curve in GraphPad Prism.

### Cytotoxicity and cell viability assay

Cytotoxicity test was conducted using Neutral Red Uptake assay ^78^. In brief, iPSC-Ns seeded in 96 well plates (1 × 10^5^ cells/well) were treated dose-dependently for 5 days. Cells were washed with PBS containing CaCl_2_ and MgCl_2_ followed by the addition of Neutral red solution (40 μg/ml in PBS, Sigma-Aldrich) and incubation at 37°C for 3 hours. Cells were then washed with PBS containing CaCl_2_ and MgCl_2_ two times and de-stained using with Neutral red de-staining solution on a plate shaker for 30 minutes, followed by reading at 540 nm with a Microplate reader. Cell viability was determined using the CellTiter-Glo® Luminescent Cell Viability Assay (Promega) following the manufacturer’s instructions. In brief, HEK cells were plated at 5 × 10^4^ cells/well, iPSC-Ns at 1 × 10^5^ cells/well in 96-well plates and treated with drugs at 10 µM for 48 hours.

### Antibodies

The following antibodies were used for western blot: anti-beta-Amyloid, 1-16 (6E10, BioLegend, 803001), anti-beta-Amyloid, 1-42 (12F4, BioLegend, 805501), β-Amyloid (Cell Signaling Technology, 8243), TUBB3 (Proteintech, 66375-1-Ig), Biotin-anti-beta-Amyloid, 17-24 (4G8, BioLegend, 800704), Anti-Amyloid Precursor Protein, C-Terminal (Sigma-Aldrich, A8717), Anti-Human sAPPbeta (Tecan (IBL), JP18957), Presenilin 1 (Cell Signaling Technology, 5643), Presenilin 2 (Cell Signaling Technology, 9979), BACE1 (Abcam, ab2077), ADAM10 (Cell Signaling Technology, 14194), Nicastrin (Cell Signaling Technology, 5665), GGA1 (Proteintech, 25674-1-AP), SORLA (Proteintech, 22592-1-AP), CD10/Neprilysin (Cell Signaling Technology, 65534), CD107a (LAMP-1) (BioLegend, 328601; Fisher Scientific, PA1-654A), SOX2 (Proteintech, 11064-1-AP), His•Tag® (Millipore, 70796-3), anti-APP C-Terminal Fragment (BioLegend, 802803). For Immunofluorescence: polyclonal Rabbit anti-CAPRIN1 (15112-1-AP, Proteintech), monoclonal mouse anti-APP (22C11, MAB348, Millipore sigma), monoclonal mouse anti-MAP2 (AP18, MA5-12826, Thermo Fisher Scientific), polyclonal rabbit anti-NANOG (PA1-097); polyclonal rabbit anti-Calnexin (10427-2-AP, Proteintech), polyclonal rabbit anti-ERGIC-53 (13364-1-AP, Proteintech), polyclonal rabbit anti-SEC31A (17913-1-AP, Proteintech), polyclonal rabbit anti-TMED9 (21620-1-AP, Proteintech), monoclonal rabbit anti-GM130 (D6B1, 12480S, Cell Signaling Technology), polyclonal rabbit anti-TGN46 (13573-1-AP, Proteintech), monoclonal rabbit anti-EEA1 (C45B10, 3288S, Cell Signaling Technology), monoclonal rabbit anti-Rab7 (D95F2, 9367S, Cell Signaling Technology); monoclonal rabbit anti-LAMP1 (D2D11, 9091S, Cell Signaling Technology). The secondary antibodies for immunofluorescence: Alexa Fluor 488® Goat anti-Mouse IgG; Alexa Fluor 488® Goat anti-Rabbit IgG; Alexa Fluor 555® Goat anti-Mouse IgG; Alexa Fluor 555® Goat anti-Rabbit IgG.

### Western blot analysis and immunoprecipitation

Cells were lysed in 1% triton X-100 buffer (50 mM Tris (pH 7.4), 150 mM NaCl, 10% glycerol, 1% Triton X-100, 1 mM EDTA) or RIPA buffer (50 mM Tris (pH7.5) 150 mM NaCl, 0.5% NP40, 0.5% deoxycholate, 0.1% SDS) supplemented with 1 mM PMSF, protease and phosphatase inhibitors. For immunoprecipitation, cell lysates were incubated with the primary antibodies (1-2 µg) overnight at 4°C and mixed with protein G beads for 4 hours. The immunoprecipitants were washed with 1% triton X-100 buffer 3 times and eluted with Laemmli buffer, followed by western blot. Western blot bands were quantified using Fiji (ImageJ) with background subtraction and normalized to loading controls.

### Real-time PCR

Total RNA was isolated using the RNeasy kit (Qiagen), followed by synthesis for cDNA with a reverse transcription reaction (Quantitect® Reverse Transcription kit). The sequences for APP and GAPDH are in Supplementary Table 1. Quantitative PCR was performed on QuantStudio 6 Flex (Applied Biosystems). The expression levels of APP were normalized to the GAPDH mRNA levels and represented relative to the expression in control cells. The results were analyzed with the delta-delta-Ct methods. Ct values were calculated using QuantStudio Design & Analysis Software, and relative expression levels were analyzed with GraphPad Prism.

### Enzyme-linked immunosorbent assay (ELISA)

iPSC-Ns or HEK-APP695^WT^ were treated with drugs, and secreted Aβ42 peptides inculture soup were quantified using the Human Amyloid β1-42 human ELISA kit (Thermo Fisher) according to the manufacturer’s protocol. Mouse brain extraction was performed as previously described^79^, with minor modifications. Mouse brain tissues were homogenized at a 1:10 weight-to-volume ratio in Tris-buffered saline (TBS; 50 mM Tris, pH 7.4, 150 mM NaCl) supplemented with 2 mM EDTA and protease inhibitors. TBS-homogenized brains were centrifuged for 1 hour at 16,000 g. To extract total Aβ, including insoluble fractions, formic acid was added to a final concentration of 70%, and the samples were thoroughly mixed by pipetting. Homogenates were centrifuged at 100,000 g for 1 hour at 4 °C, and the resulting supernatants were collected. Prior to quantification, formic acid-containing samples were neutralized with 1 M Tris base (pH 11). Aβ40 and Aβ42 levels were measured using human-specific ELISA kits (Invitrogen) and normalized to total protein concentrations determined by the BCA assay (Thermo Scientific).

### Immunofluorescence staining

Cells were grown on glass cover slips coated with 1% Matrigel and treated with drugs. Cells were washed, fixed, permeabilized, blocked, and incubated in primary antibodies diluted in blocking solution (1% v/v BSA solution/5% Goat serum in PBS containing 0.1% Triton X-100) overnight at 4°C. The cells were washed three times with PBS and incubated in secondary antibodies in a blocking solution for 1 hour at room temperature, then washed with PBS, followed by HOECHST staining (Invitrogen) for 5 minutes, and washed with PBS. The glass coverslips were put on a slide-in mounting medium (VECTASHIELD), and the edges were sealed with coverslip sealant (BIOTIUM). Image data was captured and processed with a confocal microscope system (Olympus Fluoview FV1000 or Leica DIVE) and ImageJ program.

For 3D organoids staining, organoids were fixed in 10% formalin overnight at room temperature followed by incubation in 30% sucrose solution overnight at room temperature, embedded in optimal cutting temperature compound (OCT, FisherScientific) and kept at -80°C overnight. Frozen samples were sectioned at 10 µm using a cryostat (Leica) and mounted on microscope slides (Fisherbrand™ superfrost plus, FisherScientific). Before immunolabeling, sections were rinsed in PBS for 5 minutes, permeabilized for 30 minutes in PBS containing 0.3% Triton-X 100, blocked for 1 hour in PBS containing 5% BSA and 0.1% Triton-X 100, and incubated with primary antibodies diluted in the same blocking solution. Then, secondary staining, mounting, and imaging were followed.

### Immunohistochemistry of 3D organoids, human AD patient brain samples, and mouse Brain samples

Autopsy samples of brain tissues were provided and immunohistochemistry was performed in the Department of Pathology and Laboratory Medicine, Indiana University School of Medicine following the institutional protocols. Paraffin-embedded 3D organoids and human AD patient brain samples subjected to Hematoxylin and Eosin (H&E), synapses, GFAP, CAPRIN1 (polyclonal Rabbit anti-CAPRIN1, 15112-1-AP, Proteintech), APP (monoclonal mouse anti-APP/Aβ (6E10), 803001, Biolegend; MAB348, Millipore sigma) in a Dako automated instrument using the Dako Flex detection system. The primary antibodies and secondary antibodies were used in colocalization experiments with human AD brain biopsy samples, then imaged using a confocal microscope system.

Thioflavin-S was stained with deparaffinization and rehydration of mouse brain sections using a series of xylene and ethanol washes. The sections were then incubated in filtered 1% aqueous Thioflavin-S for 8 minutes at room temperature, protected from light, followed by sequential washes in ethanol and distilled water. Finally, slides were covered with aqueous mounting solution, dried overnight in the dark, sealed with nail polish, and stored at 4°C to prevent fading before analysis.

### Imaging Analysis

ImageJ (Fiji) program (NIH, Bethesda, MD, USA) was used to analyze images ^80^. Pearson’s correlation coefficients were calculated with JACoP plugin from two channels (BIOP). For colocalization/trafficking analysis with organelles in iPSC-Ns, HEK293 cells, and 3D organoids, APP or CAPRIN1 with organelles signals were segmented using a trainable weka segmentation plugin in Fiji as previously described^81^. Colocalized areas were calculated in the image calculator tool (‘Or’ function) to get an overlayed area for the segmented signals, followed by using the analyze particle tool in Fiji to determine by > 1-pixel overlapping of the segmented signal. To determine the fluorescence intensity, the corrected total cell fluorescence (CTCF) was used using the formula: CTCF = Integrated Density – (Area of selected cell x Mean fluorescence of background readings), as described previously^82^.

Bright-field images of immunoperoxidase-stained and Thioflavin S-stained brain sections were exported as 16-bit RGB TIFF files and analyzed using ImageJ (Fiji), following previously established protocols ^83, 84^. To isolate the DAB signal for quantification, color deconvolution was performed using the “H DAB” setting in the Colour Deconvolution plugin in ImageJ. The resulting “Colour_2” image, corresponding to the DAB chromogen, was converted to 8-bit grayscale for further analysis. Manual thresholding was applied to the DAB channel using the threshold adjustment tool in ImageJ. Threshold levels were individually adjusted for each image to match the visual intensity of the stained signal, and the thresholded image was displayed in black and white mode for clarity. Regions of interest (ROIs), specifically the subiculum, hippocampus, and cortex, were manually delineated based on anatomical landmarks and saved using the ROI Manager. For each ROI, the DAB-positive area was quantified using the Analyze Particles plugin with default particle size settings. The percentage of DAB-positive area was calculated as the ratio of the DAB-positive area to the total area of each ROI.

### Recombinant protein purification from E. coli

For full-length CAPRIN1 purification, pET-6HIS-TEV-hCAPRIN1 (Vectorbuilder) were transformed into E. coli BL21 (DE3) followed by culturing in Luria broth (LB) with 100 µg/mL of Ampicillin overnight at 37°C. 10 ml of primary culture was inoculated into fresh LB medium with Ampicillin, cultured until OD_600_ reached 0.8, and expression of CAPRIN1 was induced with 0.5 mM isopropyl-β-D-thiogalactoside at 37°C for 4 hours. After centrifugation at 3,000 rpm for 20 minutes at 4°C, pellets were resuspended with prechilled lysis buffer (50 mM Na_2_HPO_4_, 300 mM NaCl, 10% Glycerol, 0.1% Triton X-100, 10 mM Imidazole at pH 8.0), followed by sonication for 16 minutes (10 seconds on and 30 seconds off for 24 cycles) on ice and centrifuged for 40 minutes (40,000 rpm at 4°C). The supernatant was mixed with Ni-NTA agarose beads (Qiagen) overnight at 4°C, then loaded on a PD-10 column (GE healthcare). The columns were washed with buffer (50 mM Na_2_HPO_4_, 300 mM NaCl, 10% Glycerol, 10 mM Imidazole at pH 8.0) and eluted with elution buffer (50 mM Na_2_HPO_4_, 300 mM NaCl, 10% Glycerol, 250 mM Imidazole at pH 8.0), followed by dialysis (50 mM Na_2_HPO_4_, 250 mM NaCl, 10% Glycerol at pH 8.0) for 2 hours at 4°C. The samples were concentrated using Millipore Amicon filters (Millipore) and dialyzed with storage buffer (50 mM Na_2_HPO_4_, 200 mM NaCl, 50% Glycerol at pH 8.0) for storage at -80°C.

### Recombinant protein from insect cells

The DNA cassette (10xHis-3C site or 10xHis-MBP-3C site) was inserted immediately 5’ of the *Bam*HI site in the pFastBac vector (Invitrogen, 10360014) using the SLIC method ^85^, yielding constructs termed pSEP0(10His-3C) (pYT1353), or pSEP8 (10His-MBP-3C) (pYT1566) vector, respectively. The open reading frame (ORF) of human CAPRIN1 was codon-optimized, synthesized, and sub-cloned into the BamHI and HindIII site of a pSEP0 vector, yielding pSEP0-CAPRIN1 (10xHis-3C-CAPRIN1) (pYT1837) vector. The open reading frame (ORF) of a full-length human APP (APP695) was codon-optimized, synthesized, and sub-cloned into the BamHI and HindIII site of a pSEP8 vector, yielding pSEP8-APP695 (10xHis-MBP-3C-APP695) (pYT1842) vector. Generation of baculovirus expressing 10xHis-CAPRIN1 or 10xHis-MBP-APP695 using pSEP0-CAPRIN1 or pSEP8-APP695 vector in Sf9 cells (Expression Systems, Inc) was performed as described previously ^86^. Expression of recombinant CAPRIN1, or APP695 was optimized using the TEQC method ^86^ in which a 200 ml culture of Hi5 cells (Expression Systems, Inc, 94-002S) was infected at an estimated multiplicity of infection (eMOI) of 4. After a 96-hour incubation at 27°C, cells were harvested, frozen in liquid nitrogen and kept at -80°C until use.

For the purification of CAPRIN1, cell pellet was lysed in 50 mM Hepes-KOH pH 7.6, 400 mM potassium chloride (KCl), 10 % Glycerol, 5 mM β-mercaptoethanol, 10 mM Imidazole, and 0.5 ml of 100X protease cocktail (6 mM Leupeptin, 0.2 mM Pepstatin A, 20 mM Benzamidine, and 10 mM PMSF). Cell lysate was stirred for 30 min followed by centrifugation at 100,000 g for 45 min. The supernatant was loaded onto Ni-resin (HIS-Select, Sigma-Aldrich) and washed with buffer 1 (50 mM Hepes-KOH pH7.6, 1 M KCl, 5% Glycerol, 10 mM Imidazole, and 5 mM β-mercaptoethanol), and buffer 2 (50 mM Hepes-KOH pH7.6, 2 M KCl, 5% Glycerol, 10 mM Imidazole, and 5 mM β-mercaptoethanol) followed by lysis buffer and the low salt buffer (50 mM Hepes-KOH pH7.6, 200 mM KCl, 5% Glycerol, 10 mM Imidazole, and 5 mM β-mercaptoethanol). 10xHis-CAPRIN1 was eluted from Ni-resin with the elution buffer (50 mM Hepes-KOH pH7.6, 200 mM potassium acetate, 5% Glycerol, 300 mM Imidazole, and 5 mM β-mercaptoethanol). Elution fractions containing 10xHis-CAPRIN1 were pooled and dialyzed overnight against the low salt buffer. Dialysate was concentrated by spin column (Amicon 10k cut-off) to 4.0 mg/ml. Recombinant CAPRIN1 was further purified by 10–30% (v/v) glycerol gradient containing the buffer (50 mM Hepes-KOH pH7.6, 200 mM KCl, 5% glycerol and 5 mM β-mercaptoethanol). After centrifugation at 35,000 rpm in a Beckman SW Ti-40 rotor for 19 hours at 4°C, the gradient was fractionated using a PGF Piston Gradient Fractionator (BioComp Instruments). Fractions containing 10xHis-CAPRIN1 were pooled and dialyzed overnight against the low salt buffer. Dialysate was concentrated by spin column (Amicon 10k cut-off) to 4 mg/ml. Concentration of purified protein was measured by Bradford method. When purifying APP695, the same procedure as the CAPRIN1 purification was used. Buffer 1 (50 mM Hepes-KOH pH7.6, 1 M KCl, 5% Glycerol, 5 mM β-mercaptoethanol, and 0.1% DDM), wash buffer 2 (50 mM Hepes-KOH pH7.6, 2 M KCl, 5% Glycerol, 5 mM β-mercaptoethanol, and 0.1% DDM), low salt buffer with 0.1% DDM, elution buffer (50 mM Hepes-KOH pH7.6, 200 mM potassium acetate, 5% Glycerol, 50 mM Maltose, and 5 mM β-mercaptoethanol), and Amylose-resin were used.

### CAPRIN1-APP interaction by glycerol gradient

About 1 mg of 10xHis-MBP-APP695 was mixed with ∼1 mg of 10xHis-CAPRIN1 and loaded onto 10-30% (v/v) glycerol gradient containing the buffer (50 mM Hepes-KOH pH7.6, 200 mM KCl, 5 mM β-mercaptoethanol, and 0.1% DDM). For controls, ∼1 mg of 10xHis-MBP-APP695 (full-length) alone was loaded onto 10-30% (v/v) glycerol gradient as well. After centrifugation at 35,000 rpm in a Beckman SW Ti-40 rotor for 19 hours at 4°C, the gradient was fractionated as described above. For fractions containing 10xHis-CAPRIN1, a PGF Piston Gradient Fractionator was used. Fractions containing 10xHis-CAPRIN1, 10xHis-MBP-APP695 alone or mixture were subjected to 4-12% NuPAGE (Invitrogen) and stained with Coomassie blue.

### *In vitro* binding assay

When CAPRIN1 was used as a bait, recombinant CAPRIN1 (200 ng) was mixed with anti-CAPRIN1 antibodies (1µg) in IP buffer (25 mM Tris, 150 mM NaCl, 5% Glycerol, and 1% Triton X-100 at pH 7.4) overnight at 4°C, followed by the addition of Protein-G agarose beads for 4 hours at 4°C. CAPRIN1 conjugated beads were then washed with binding buffer (20 mM HEPES, 100 mM NaCl, 0.01% Triron X-100, and 5% Glycerol at pH 7.25) before the addition of recombinant human APP (prey) for 1 hour at 4°C. Beads were washed and eluted with Laemmli buffer with boiling, followed by Western blotting.

When APP was used as a bait, biotin-conjugated APP antibody (1 µg; 4G8, Biolegend) was pre-mixed with recombinant APP (100 ng, Biolegend) in pulldown buffer (0.1% NP-40, 50 mM Tris, 150 mM NaCl, 0.2 mM EDTA, 100 mM KCl, 1 mM MgCl_2,_ 0.2 mM CaCl_2,_ and 10% glycerol) overnight at 4°C, followed by addition of 0.5 mg of Streptavidin magnetic beads (Pierce) for 1 hour at 4°C. After washing the beads, recombinant human CAPRIN1 (full length, 100 ng) was added together with dose-dependent 0043 in the binding buffer for 1 hour at 4°C. Beads were washed and eluted with Laemmli buffer with boiling, followed by Western blotting.

### Surface Plasmon Resonance (SPR)

SPR experiments were conducted to investigate the interaction between APP695 and CAPRIN1 proteins using a Biacore T200 instrument (Cytiva). Both proteins were expressed in Tni (Hi5) insect cells and purified to homogeneity via tandem affinity and ion-exchange. Protein purity and integrity were confirmed by SDS-PAGE and size-exclusion chromatography prior to analysis. All experiments were performed in a running buffer consisting of 50 mM HEPES at pH7.0, 100 mM KCl, 10% glycerol, and 0.01% (v/v) Tween-20 supplemented with 0.1% (w/v) Sarkosyl (pH 7.4). APP695 was serially diluted in running buffer to final concentrations of 15 µM, 5 µM, 1 µM, and 0.3 µM.

#### Chip Preparation

A CM5 sensor chip (Cytiva) was activated using a 1:1 (v/v) mixture of 0.1 M N-hydroxysuccinimide (NHS) and 0.1 M N-ethyl-N′-(3-dimethylaminopropyl)carbodiimide (EDC) to generate reactive ester groups. CAPRIN1 was diluted to 50 µg/mL in 10 mM Hepes-KOH buffer (pH 6.0) and immobilized on the chip surface via standard amine coupling to achieve immobilization. Unreacted sites were blocked with 1 M ethanolamine-HCl (pH 8.5). A reference flow cell was prepared similarly but without protein immobilization to control for non-specific binding and bulk refractive index changes.

#### Binding Assay

Binding interactions were measured at 25°C with a flow rate of 30 µL/min. APP695 solutions were injected over the CAPRIN1-immobilized and reference flow cells for 45 s (association phase), followed by a 240 s dissociation phase with running buffer alone. Internal control injections for one concentration of APP695 were performed to ensure reproducibility for every round of the experiment. 0043 and 0152 produced no noticeable interaction with CAPRIN1 alone. APP(1–300) was also tested for binding to CAPRIN1 under identical conditions. To assess potential non-specific interactions, 0043 and 0152 were tested for binding to APP-695 captured on a CM5 chip using a steady-state model.

#### Data Analysis

Sensograms were processed using Biacore T200 Evaluation Software. Signals from the reference flow cell were subtracted from those of the CAPRIN1 flow cell to account for non-specific binding and bulk refractive index effects. Baseline drift was corrected by subtracting responses from blank buffer injections. Binding data were globally fitted to a heterogeneous ligand model to calculate kinetic parameters, including the association rate constant (k_on_) and dissociation rate constant (k_off_). Equilibrium dissociation constants (K_D_) were derived from the ratio k_off_/k_on_.

### Hydrogen-Deuterium Exchange Mass Spectroscopy (HDX-MS)

#### Protein Preparation

APP695 and CAPRIN1 proteins were buffer exchanged into HDX-compatible buffer (10 mM HEPES, 150 mM NaCl, pH 7.4) supplemented with 0.1% (w/v) Sarkosyl (pH 7.4) using dialysis and subsequently diluted to 10 µM. For the small molecule binding experiment, 0152 (dissolved in DMSO) was diluted to 1 mM in HDX-compatible buffer and then subsequently diluted in the complex to maintain a final DMSO concentration of ∼1%.

#### Complex Formation and HDX Reaction Setup

APP695 and CAPRIN1 proteins were mixed in equimolar ratios (10 µM each) and incubated on ice for 60 minutes to allow complex formation. HDX reactions were initiated by diluting the protein complex 1:10 into D_2_O buffer (10 mM HEPES, 150 mM NaCl, pH 7.4, prepared with 99.9% D_2_O) at 0°C. The exchange was allowed to proceed for varying times (1 min and 10 min) before quenching with ice-cold quench buffer (2M Urea, 0.8% Formic Acid, 20 mM TCEP). HDX reactions were performed in triplicates for individual data points.

#### Digestion and Peptide Analysis

Vanquish (ThermoFisher) with QE HF was used for LC-MS analysis. Samples were manually injected on a 100ul sample loop. The sample was then passed through a protease type XIII/pepsin column (NovaBioassays) at 100ul/min with 0.2% formic acid and trapped on a self-packed Poros R10, 2.1X4cm trap column for 4 min. Peptides were separated on a 2.1mmx5cm C18 column (1.9 u, Hypersil Gold, Thermofisher) with a multistep gradient: Gradient: 0-4 min: 2%B, from 2% B to 10% B in 1.3 min, 10 to 40%B in 9.7 min, and 40-85%B in 0.7 min. The digestion and separation were conducted at 5C. The mass was measured with a resolution of 120,000 and a mass range from 300-2000 with top20 peaks fragmented by HCD (relative collision energy 27) and scanned with a resolution 15,000 with a dynamic exclusion for 20 sec.

#### Data analysis

The LC-MSMS data were analyzed using Proteome Discoverer 3.0 with Sequest HT against a custom database composed of target proteins and kinases. MS window was set at +/- 10 ppm, MS/MS window was set at +/- 0.02Da. N-terminal acetylation was added as dynamic protein terminus modification, Oxidation at methionine, and phosphorylation at serine/threonine/tyrosine were set as dynamic modifications. Protease was set as non-specific. The results were validated by Fix Value PSM Validator with max Delta CN < 0.05. PhosphoRS was used to calculate phosphorylation site confidence. If a site was 90% or more phosphorylated, only phosphorylated forms were considered for HDX analysis. The HDX time point results were analyzed using HDexaminer 3.4 (Sierra Analytics) with manual inspection.

### HDX-MS with 0152

#### Complex Preparation and HDX Reaction Setup

APP695 and CAPRIN1 proteins were pre-incubated in equimolar concentrations to form a stable complex as previously described. 0152 was added to the complex at a final concentration of 100 µM (10:1 molar ratio of ligand to protein complex) and incubated for an additional 60 minutes at 4°C. HDX was initiated as described above, with time points at 1 min and 10 min. Samples were quenched and processed identically to the protein-protein interaction experiment. Each HDX-MS experiment was performed in triplicate to ensure reproducibility. Controls included individual proteins, the protein-protein complex without 0152, and the complex with DMSO (vehicle control) to account for solvent effects.

#### Data Analysis

Deuterium incorporation was compared between the protein complex in the presence and absence of 0152. Changes in deuterium uptake (ΔHDX) greater than 0.5 Da with statistical significance (p < 0.01) were mapped onto the protein structures. Regions exhibiting reduced deuterium uptake upon 0152 binding were identified as the small-molecule binding site.

##### Synthesis of 0043

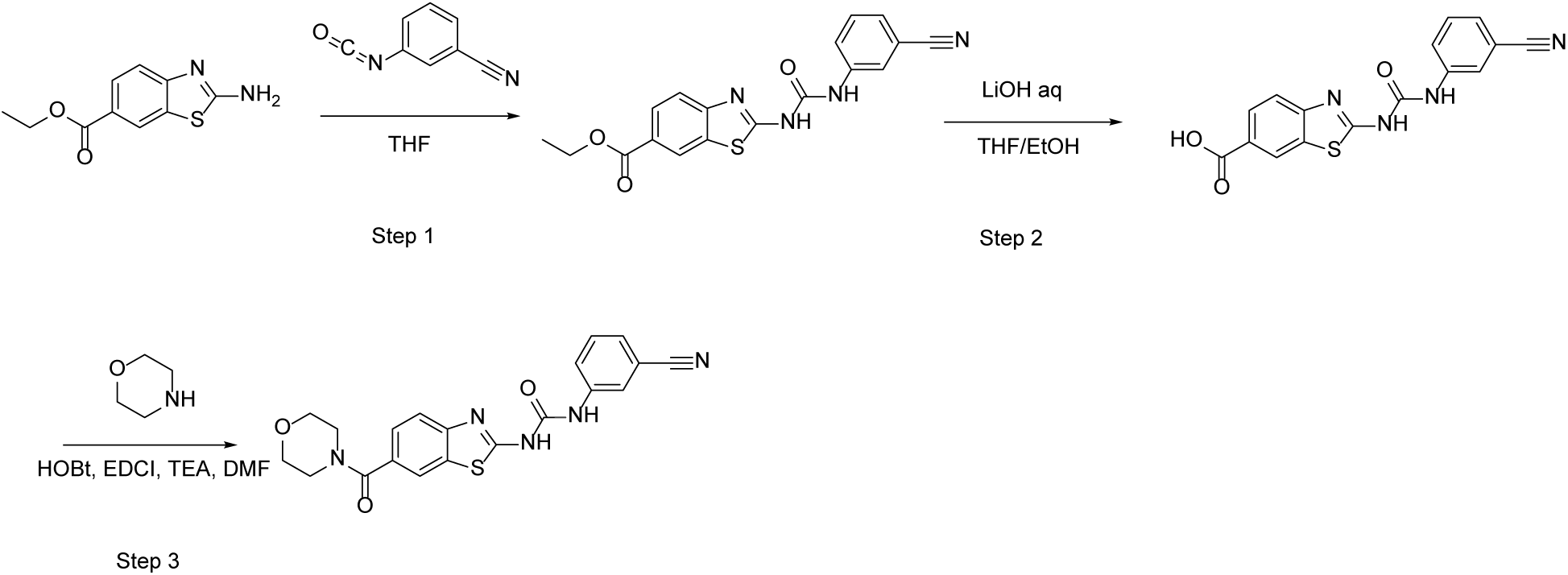

#### Step 1. Synthesis of methyl 2-[(3-cyanophenyl)carbamoylamino]-1,3-benzothiazole-6-carboxylate

Methyl 2-amino-1,3-benzthiazole-6-carboxylate (2 g, 9.6 mmol) was suspended in Tetrahydrofuran (60 mL) and reacted with 3-isocyanatobenzonitrile (1.65 g, 11.5 mmol) at 25℃for 16 hours. TLC was run in 1:1 Ethyl acetate/Hexane to show completion. The reaction mixture was concentrated then diluted with 40 mL of ethanol. After heating the reaction mixture to 70℃, 40 mL of dichloromethane was added and the reaction was stirred for 30 minutes. The solid was then cooled to 25℃, filtered, and rinsed with ethyl alcohol to give the title compound as a white solid (2.8 g, yield: 83%).

#### Step 2. Synthesis of 2-[(3-cyanophenyl)carbamoylamino]-1,3-benzothiazole-6-carboxylic acid

Methyl 2-[(3-cyanophenyl)carbamoylamino]-1,3-benzothiazole-6-carboxylate (2.8 g, 7.9 mmol) was suspended in a 2:1 solution of Tetrahydrofuran / Ethyl alcohol (80 mL). An aqueous 1N Lithium hydrate solution (58.1 mmol) was added to the reaction and stirred at 25℃ for 2 days. TLC in 1:1 Ethyl acetate/Hexane showed completion. The reaction was concentrated to remove the organic solvents, and the residue was diluted with 20 mL of H_2_O and 80 mL of Isopropyl alcohol. The pH was adjusted to 5 using Acetic acid and after stirring for 1 hour, the reaction was filtered and rinsed with Isopropyl alcohol to obtain the title compound as a white solid (2.3 g, yield: 88%).

#### Step 3. Synthesis of 0043

2-[(3-cyanophenyl)carbamoylamino]-1,3-benzothiazole-6-carboxylic acid (2.3 g, 6.8 mmol) was suspended in DMF (90 mL), then EDCI (2.1 g, 13.6 mmol) and HOBt (1.1 g, 8.16 mmol) were added to the reaction. After stirring at 25℃for 10 minutes, Morpholine (2.9 g, 34 mmol) and Triethylamine (9.4 mL, 68 mmol) was added, then the reaction for stirred for 2 days at 25℃. TLC in 1:1 Ethyl acetate/Hexane showed completion, the reaction was poured into 200 mL of H_2_O and washed three times with 200 mL of Ethyl acetate. The organic washes were combined and concentrated to ∼100 mL, and 100 mL of H_2_O was added. The organic and aqueous were left alone for 8 hours until a white solid precipitated out of the solution. The solid was then filtered and rinsed with H_2_O to give the title compound 0043 (1.2 g, yield: 43%) as a white solid.

##### Synthesis of 0152

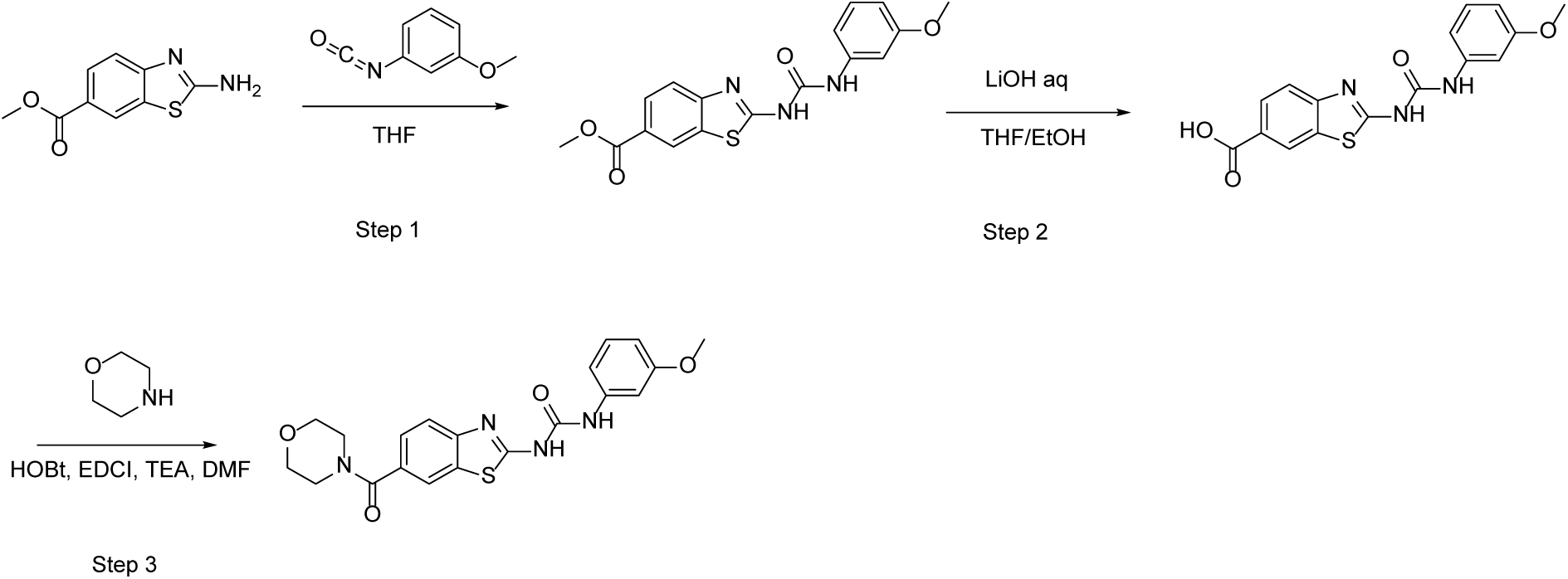

#### Step 1. Synthesis of methyl 2-(3-(3-methoxyphenyl)ureido)benzo[d]thiazole-6-carboxylate

Methyl 2-amino-1,3-benzthiazole-6-carboxylate (2 g, 9.6 mmol) was suspended in THF (60 mL) and reacted with 1-methoxy-3-isocyanato-benzene (1.72 g, 11.5 mmol) at 25℃ for 16 hours. TLC was run in 1:1 Ethyl acetate/Hexane to show completion. The reaction was concentrated then diluted with 40 mL of Ethyl alcohol. After heating the reaction mixture to 70℃, 40 mL of Dichloromethane was added and the reaction was stirred for 30 minutes. The solid was then cooled to 25℃, filtered and rinsed with Ethyl alcohol to give the title compound as a white solid (2.6 g, yield: 76%).

#### Step 2. Synthesis of 2-(3-(3-methoxyphenyl)ureido)benzo[d]thiazole-6-carboxylic acid

Methyl 2-(3-(3-methoxyphenyl)ureido)benzo[d]thiazole-6-carboxylate (2.6 g, 7.3 mmol) was suspended in a 2:1 solution of Tetrahydrofuran / Ethyl alcohol (80 mL). An aqueous 1N Lithium hydrate solution (58.1 mmol) was added to the reaction and stirred at 25℃ for 2 days. TLC in 1:1 Ethyl acetate/Hexane showed completion. The reaction was concentrated to remove the organic solvents and was diluted with 20 mL of H_2_O and 80 mL of Isopropyl alcohol. The pH was adjusted to 5 using Acetic acid, and after stirring for 1 hour, the reaction was filtered and rinsed with Isopropyl alcohol to give the title compound as a white solid (2.4 g, yield: 97%).

#### Step 3. Synthesis of 0152

2-(3-(3-methoxyphenyl)ureido)benzo[d]thiazole-6-carboxylic acid (2.4 g, 7.3 mmol) was suspended in DMF (90 mL), then EDCI (2.8 g, 14.6 mmol) and HOBt (1.19 g, 8.78 mmol) was added to the reaction. After stirring at 25℃ for 10 minutes, morpholine (2.55 g, 29.3 mmol) and triethylamine (8.21 mL, 58.5 mmol) were added, then the reaction for stirred for 2 days at 25℃. TLC in 1:1 Ethyl acetate/Hexane showed completion, the reaction was poured into 200 mL of H_2_O and washed three times with 200 mL of Ethyl acetate. The organic washes were combined and concentrated to ∼100 mL, and 100 mL of H_2_O was added. The organic and aqueous were left alone for 8 hours until a white solid precipitated out of solution. The solid was then filtered and rinsed with H_2_O to give 0152 as a white solid (800 mg, yield: 27%).

### Statistics

#### Statistical analysis

Statistical analyses were performed using GraphPad Prism 9.0 software. Data are presented as mean ± standard deviation (SD), and the number of biological replicates (n) is indicated in each figure legend. Comparisons between two groups were analyzed using unpaired two-tailed Student’s t-tests. For comparisons among three or more groups, one-way analysis of variance (ANOVA) followed by Tukey’s multiple comparisons test was used. For experiments involving two independent variables, two-way ANOVA with appropriate post hoc tests (Tukey’s or Bonferroni’s) was applied. Dose–response curves and IC₅₀ values were generated using nonlinear regression with a three-parameter variable slope model. For thermal shift assays (CETSA and ThermoFluor), melting curves were fitted using Boltzmann’s sigmoidal function. Pearson’s correlation coefficients were calculated for confocal image colocalization analysis using the JACoP plugin in ImageJ (Fiji). Pharmacokinetic parameters were estimated using non-compartmental analysis in Phoenix WinNonlin v8.3. Statistical significance was defined as P < 0.05 and denoted as follows: P < 0.05 (*), P < 0.01 (**), P < 0.001 (***), and P < 0.0001 (****); “ns” indicates not significant.

---

**Supplementary Table 1.**
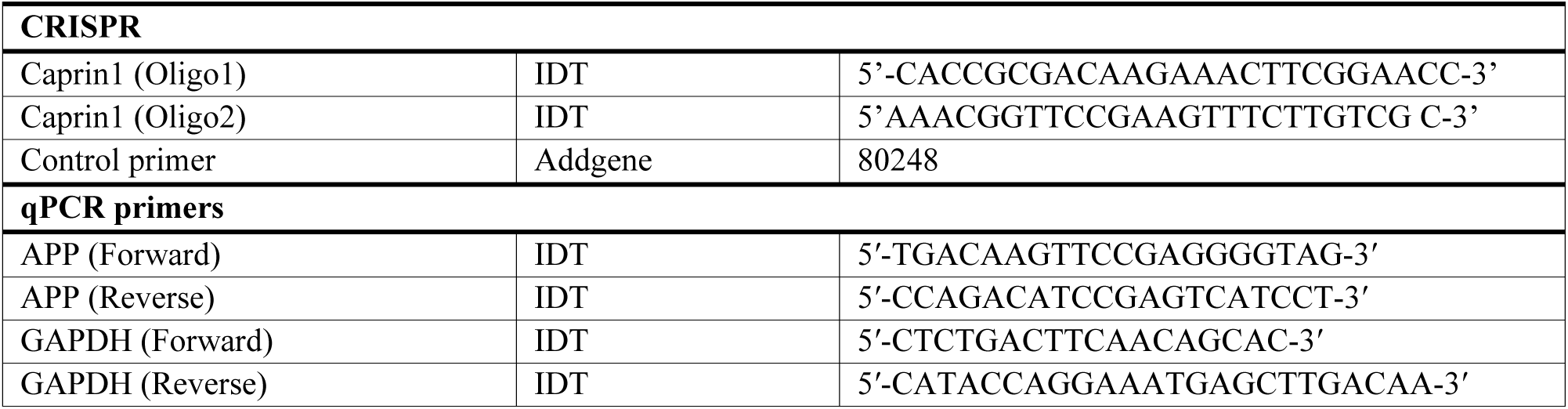
Oligonucleotides.

