## Supplementary Material for "Molecular glue degraders enhance CAPRIN1-dependent lysosomal degradation of APP in Alzheimer’s disease"

### Supplementary Figures S1-S7 and Table S1 for

#### Molecular glue degraders enhance CAPRIN1-dependent lysosomal degradation of APP in Alzheimer's disease

Sunghan Jung, Raktim Roy, Xu Wang, Bin Liu, Nancy Jaiswal, Ratan K. Rai, Yuichiro Takagi, Cen Gao, Lifan Zeng, Ho-Yin Lo, Reagan K. Wohlford, Ryan K. Higgins, Anantha Shekhar, Anita C. Bellail, Chunhai Hao

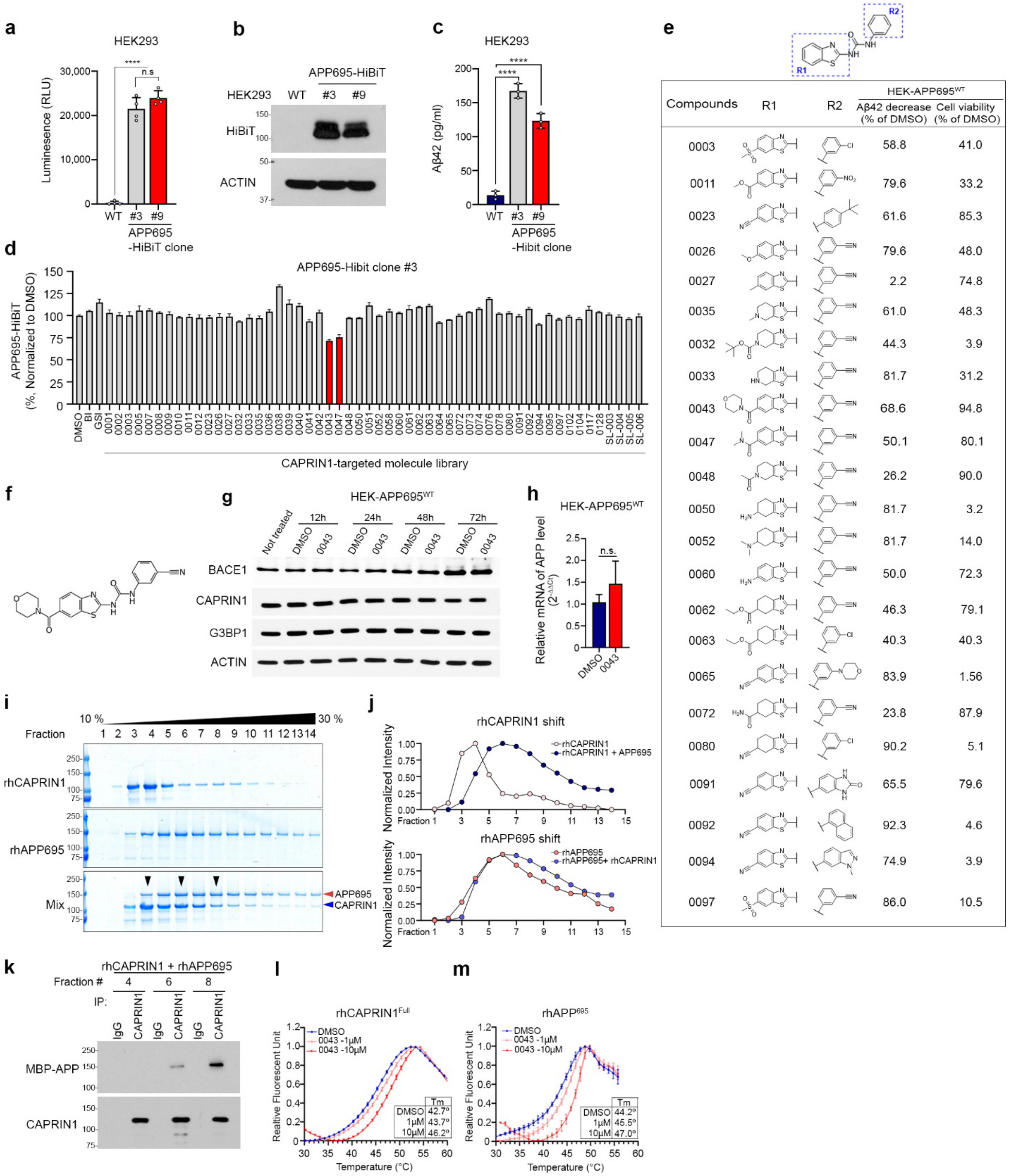
a
HEK293
Luminesence (RLU)
WT
#3
#9
APP695 -HiBiT clone
n.s
b
APP695-HiBiT
HEK293
WT
#3
#9
HiBiT
ACTIN
c
HEK293
Aβ42 (pg/ml)
APP695 -Hibit clone
d
APP695-Hibit clone #3
APP695-HiBiT (%, Normalized to DMSO)
CAPRIN1-targeted molecule library
e
R1
R2
Compounds
R1
R2
HEK-APP695WT
Aβ42 decrease (% of DMSO)
Cell viability (% of DMSO)
0003 58.8 41.0
0011 79.6 33.2
0023 61.6 85.3
0026 79.6 48.0
0027 2.2 74.8
0035 61.0 48.3
0032 44.3 3.9
0033 81.7 31.2
0043 68.6 94.8
0047 50.1 80.1
0048 26.2 90.0
0050 81.7 3.2
0052 81.7 14.0
0060 50.0 72.3
0062 46.3 79.1
0063 40.3 40.3
0065 83.9 1.56
0072 23.8 87.9
0080 90.2 5.1
0091 65.5 79.6
0092 92.3 4.6
0094 74.9 3.9
0097 86.0 10.5
f
g
HEK-APP695WT
Not treated
12h
24h
48h
72h
DMSO
0043
BACE1
CAPRIN1
G3BP1
ACTIN
h
HEK-APP695WT
Relative mRNA of APP level
n.s.
DMSO
0043
i
10 %
30 %
Fraction
rhCAPRIN1
rhAPP695
Mix
APP695
CAPRIN1
j
rhCAPRIN1 shift
rhCAPRIN1
rhCAPRIN1 + APP695
Normalized Intensity
Fraction
rhAPP695 shift
rhAPP695
rhAPP695+ rhCAPRIN1
k
rhCAPRIN1 + rhAPP695
Fraction #
IP:
IgG
CAPRIN1
MBP-APP
CAPRIN1
l
rhCAPRIN1Full
Realtive Fluorescent Unit
DMSO
0043 -1μM
0043 -10μM
Tm
DMSO 42.7º
1μM 43.7º
10μM 46.2º
Temperature (°C)
m
rhAPP695
DMSO
0043 -1μM
0043 -10μM
Tm
DMSO 44.2º
1μM 45.5º
10μM 47.0º
Temperature (°C)

**Extended Data Fig. 1 | Compound 0043 reduces APP levels in cells and promotes CAPRIN1-APP interaction in vitro.**

a, HiBiT luminescence signal in HEK293 cells stably expressing APP695-HiBiT clones (#3, #9) via lentiviral transduction (mean ± s.d., n = 4; ****p < 0.0001).

b, Western blot showing APP695-HiBiT expression in selected HEK clones (#3, #9).

c, Aβ42 levels in culture media of HEK cells expressing WT, #3, or #9 APP695–HiBiT measured by ELISA (mean ± s.d., n = 3; ****p < 0.0001).

d, Compound screening in HEK–APP695$^{WT}$ cells expressing APP695-HiBiT clone #3 from a CAPRIN1-targeted small molecule library for 24 h; changes shown as % of DMSO control, highlighting compounds 0043 and 0047 in red.

e, Chemical structures and summary table of selected compounds from the library screen, showing Aβ42 reduction and cell viability relative to DMSO.

f, Chemical structure of compound 0043.

g, Time-course western blot of BACE1, CAPRIN1, and G3BP1 in HEK-APP695WT cells treated with 0043 (10 μM) for 12, 24, 48, or 72 h.

h, qRT-PCR quantification of APP mRNA in HEK-APP695$^{WT}$ cells treated with DMSO or 0043 for 72 h (mean ± s.d., n = 3; n.s., not significant).

i, SDS-PAGE analysis of fractions from glycerol gradient centrifugation of recombinant human CAPRIN1 (rhCAPRIN1), APP695 (rhAPP695), and their mixture; Arrowheads indicate fractions showing mobility shifts in the mixture.

j, Quantification of protein distribution across gradient fractions showing elution shifts of rhCAPRIN1 (top) and rhAPP695 (bottom).

k, Immunoprecipitation (IP) of rhAPP695 and rhCAPRIN1 from size exclusion chromatography (SEC) fractions using APP or CAPRIN1 antibodies.

l,m, Thermofluor assay melt curves of recombinant full-length CAPRIN1 (l) and APP (m) pre-incubated with 0043 or DMSO; protein stability quantified by luminescence across temperature gradients (mean ± s.d., n = 3).

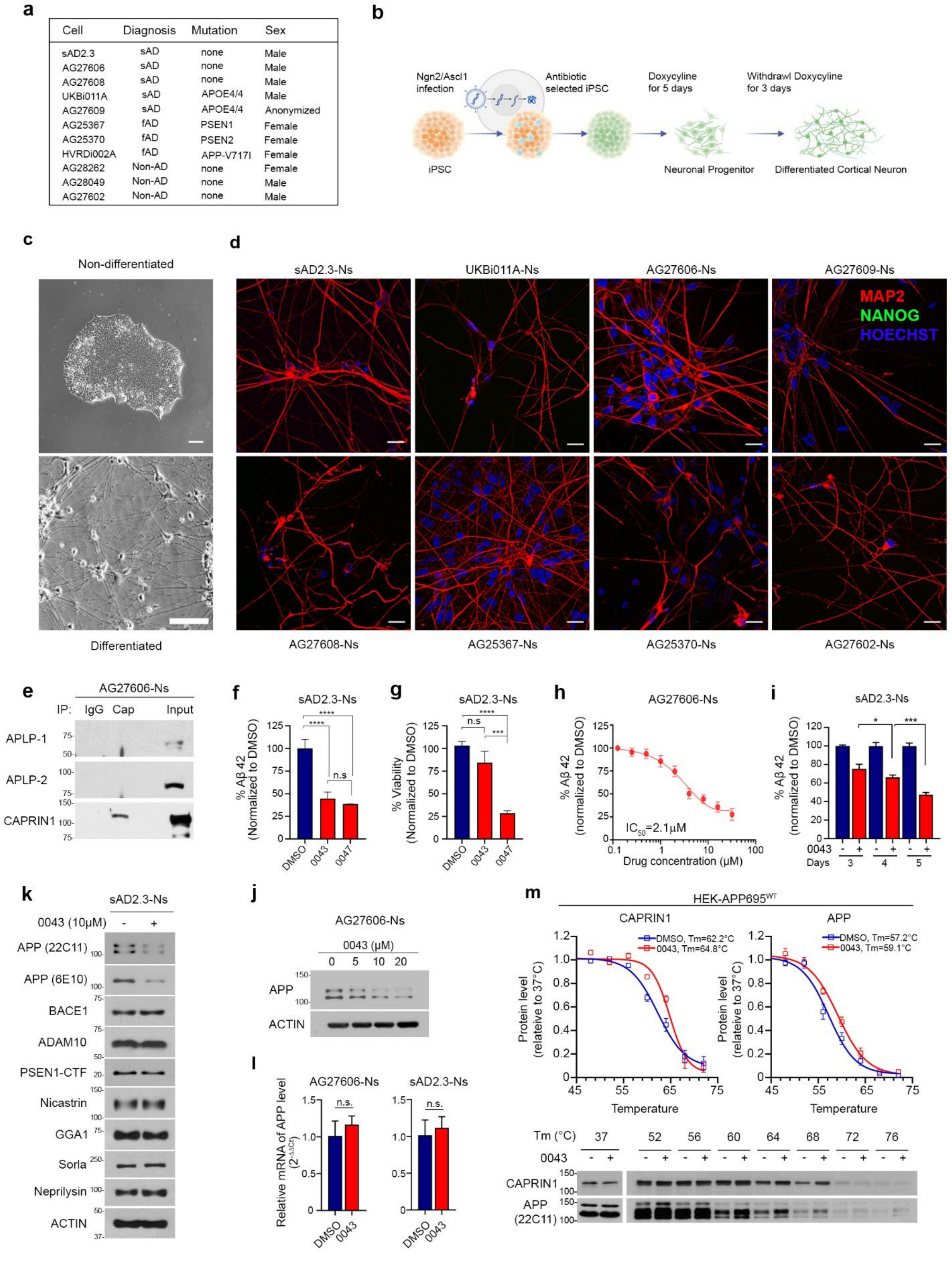
a
Cell Diagnosis Mutation Sex
sAD2.3 sAD none Male
AG27606 sAD none Male
AG27608 sAD none Male
UKBi011A sAD APOE4/4 Male
AG27609 sAD APOE4/4 Anonymized
AG25367 fAD PSEN1 Female
AG25370 fAD PSEN2 Female
HVRDi002A fAD APP-V717I Female
AG28262 Non-AD none Female
AG28049 Non-AD none Male
AG27602 Non-AD none Male
b
Ngn2/Ascl1 infection
Antibiotic selected iPSC
Doxycyline for 5 days
Withdrawl Doxycyline for 3 days
iPSC
Neuronal Progenitor
Differentiated Cortical Neuron
c
Non-differentiated
Differentiated
d
sAD2.3-Ns
UKBi011A-Ns
AG27606-Ns
AG27609-Ns
MAP2
NANOG
HOECHST
AG27608-Ns
AG25367-Ns
AG25370-Ns
AG27602-Ns
e
AG27606-Ns
IP: IgG Cap Input
APLP-1
APLP-2
CAPRIN1
f
sAD2.3-Ns
% Aβ 42 (Normalized to DMSO)
DMSO 0043 0047
n.s
g
sAD2.3-Ns
% Viability (Normalized to DMSO)
n.s
h
AG27606-Ns
% Aβ 42 (normalized to DMSO)
IC50=2.1μM
Drug concentration (μM)
i
sAD2.3-Ns
% Aβ 42 (normalized to DMSO)
0043
Days 3 4 5
k
sAD2.3-Ns
0043 (10μM) - +
APP (22C11)
APP (6E10)
BACE1
ADAM10
PSEN1-CTF
Nicastrin
GGA1
Sorla
Neprilysin
ACTIN
j
AG27606-Ns
0043 (μM)
0 5 10 20
APP
ACTIN
l
AG27606-Ns
sAD2.3-Ns
Relative mRNA of APP level (2^-ΔΔCt)
n.s.
DMSO 0043
m
HEK-APP695WT
CAPRIN1
DMSO, Tm=62.2°C
0043, Tm=64.8°C
APP
DMSO, Tm=57.2°C
0043, Tm=59.1°C
Protein level (relateive to 37°C)
Temperature
Tm (°C) 37 52 56 60 64 68 72 76
0043
CAPRIN1
APP (22C11)

**Extended Data Fig. 2 | Validation of iPSC-derived neuronal differentiation and compound 0043–mediated APP/Aβ42 regulation.**

a, Table summarizing diagnosis, genotype, and sex of iPSC lines used.

b, Schematic of Ngn2/Ascl1-based cortical neuron differentiation from iPSCs using doxycycline-inducible system.

c, Representative phase-contrast images before and after differentiation (scale bars, 100 μm).

d, Confocal images of differentiated neurons stained for MAP2 (red), NANOG (green), and Hoechst (blue) (scale bars, 20 μm).

e, CAPRIN1 IP in AG27606-Ns, followed by western blot of APLP1/2.

f, Aβ42 levels in culture media of sAD2.3 neurons treated with DMSO, 0043, or 0047 (10 μM, 72 h) by ELISA (mean ± s.d., $n = 3$; ****$P < 0.0001$; n.s., not significant).

g, Cell viability of sAD2.3 neurons after 72 h treatment with compounds measured by CellTiter-Glo (mean ± s.d., $n = 3$).

h, Dose-response curve of Aβ42 reduction in AG27606-Ns treated with 0043 for 5 days ($IC_{50}$ = 2.1 μM; mean ± s.d.).

i, Time-course analysis of Aβ42 in sAD2.3 neurons treated with 0043 (10 μM) for 3–5 days (mean ± s.d.; *$p < 0.05$, **$p < 0.01$).

j,k, Western blots of APP and related proteins in AG27606-Ns (j) and sAD2.3-Ns (k) treated with 0043 (10 μM, 72 h).

l, qRT-PCR quantification of APP mRNA in AG27606-Ns and sAD2.3 neurons treated with DMSO or 0043 for 72 h (mean ± s.d., $n = 3$; n.s., not significant).

m, Cellular thermal shift assay (CETSA) in HEK-APP695$^{WT}$ cells treated with DMSO or 0043 (10 μM, 1 h), showing thermal stability curves and representative western blots; apparent melting temperatures (Tm) indicated.

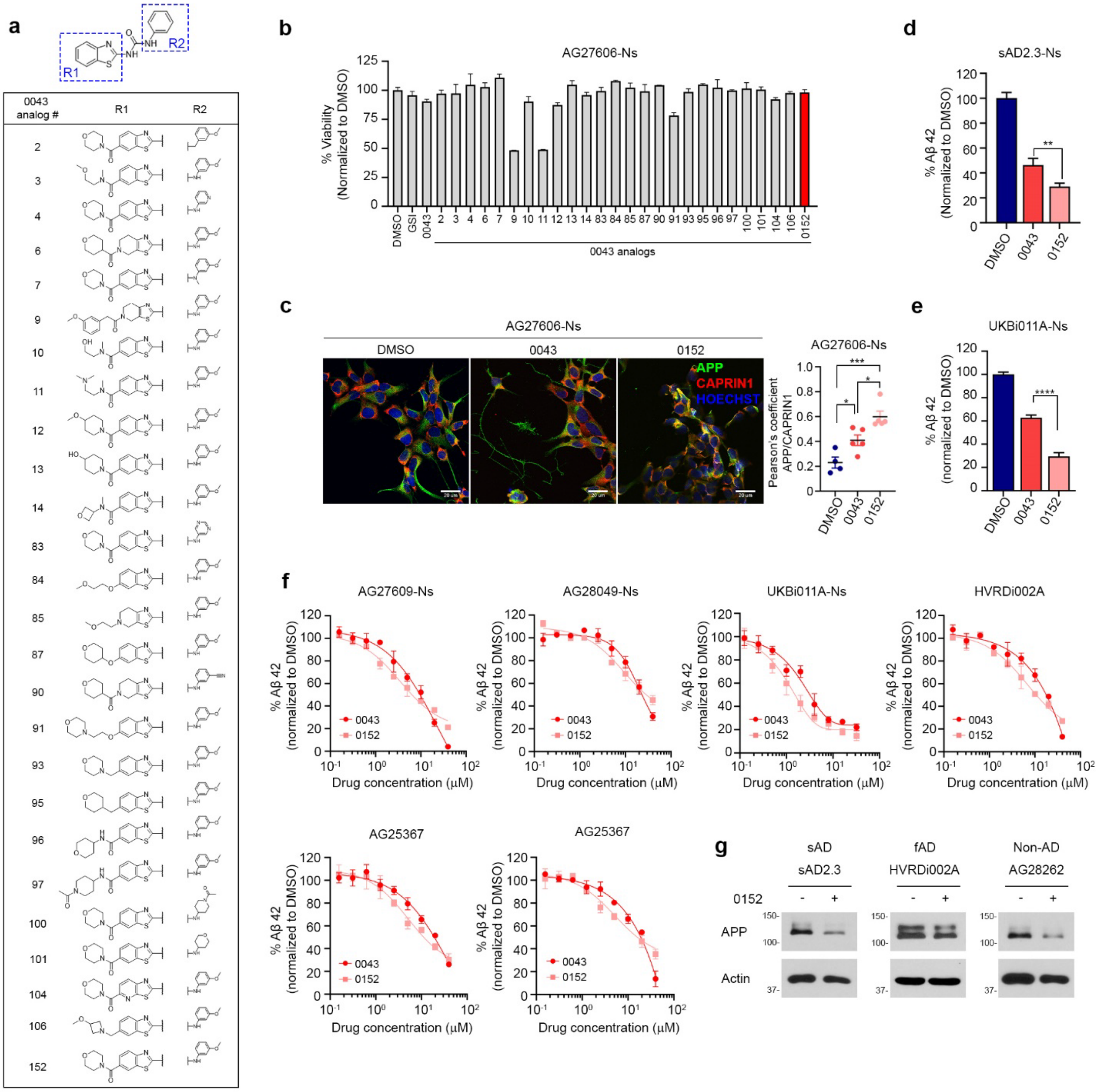
a
R1
R2
NH
0043 analog #
R1
R2
2
3
4
6
7
9
10
11
12
13
14
83
84
85
87
90
91
93
95
96
97
100
101
104
106
152
b
AG27606-Ns
% Viability (Normalized to DMSO)
DMSO
GSI
0043
0043 analogs
c
AG27606-Ns
DMSO
0043
0152
APP
CAPRIN1
HOECHST
Pearson's coefficient APP/CAPRIN1
d
sAD2.3-Ns
% Aβ 42 (Normalized to DMSO)
e
UKBi011A-Ns
% Aβ 42 (normalized to DMSO)
f
AG27609-Ns
AG28049-Ns
UKBi011A-Ns
HVRDi002A
AG25367
AG25367
% Aβ 42 (normalized to DMSO)
Drug concentration (μM)
0043
0152
g
sAD
sAD2.3
fAD
HVRDi002A
Non-AD
AG28262
0152
APP
Actin

**Extended Data Fig. 3 | 0043 analog screening and 0152 efficacy across AD and non-AD iPSC-derived neurons.**

a, Chemical structures of 0043 analogs; table summarizes structural diversity at R1 and R2 positions.

b, Cell viability of AG27606-Ns treated with 0043 and 20 analogs (10 μM, 72 h) measured by CellTiter-Glo (mean ± s.d., n = 3).

c, Confocal images of AG27606-Ns treated with DMSO, 0043, or 0152 (10 μM, 24 h), stained for APP (green), CAPRIN1 (red), and Hoechst (blue); scale bars, 20 μm. APP-LAMP1 colocalization quantified (mean ± s.d., $n \geq 4$; one-way ANOVA with Tukey's).

d,e, Aβ42 levels in culture media of sAD2.3-Ns (d) and UKBi011A-Ns (e) treated with DMSO, 0043, or 0152 (10 μM, 72 h) by ELISA (mean ± s.d., n = 3; $^{**}p < 0.01$, $^{****}p < 0.0001$).

f, Dose-response curves of Aβ42 reduction with 0043 or 0152 treatment for 5 days.

g, Western blot of APP in iPSC-derived neurons from sAD, fAD, and non-AD donors treated with 0152 (10 μM, 72 h).

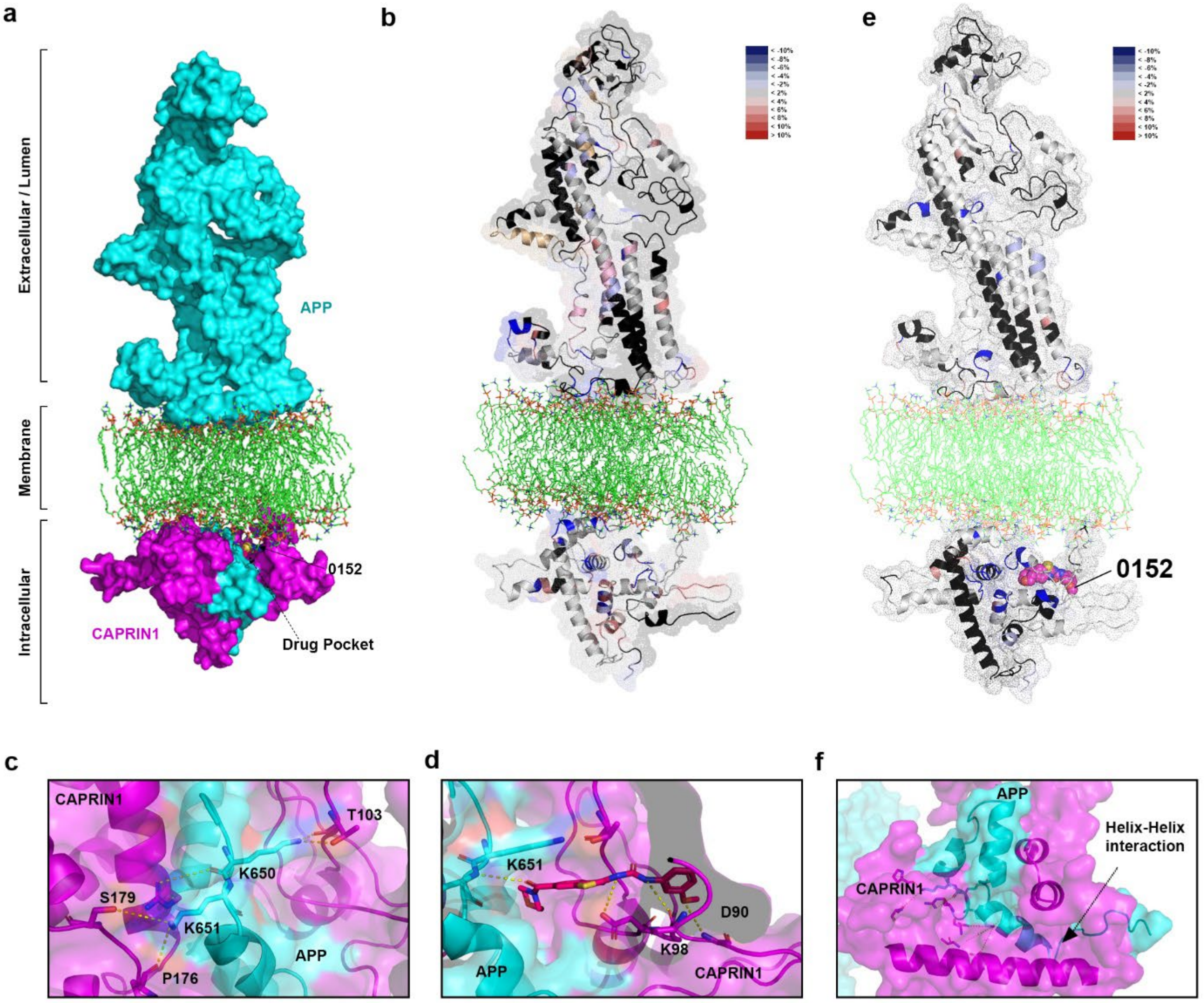
a
Extracellular / Lumen
Membrane
Intracellular
APP
CAPRIN1
0152
Drug Pocket
b
e
0152
c
CAPRIN1
T103
K650
S179
K651
P176
APP
d
K651
D90
K98
APP
CAPRIN1
f
APP
Helix-Helix
interaction
CAPRIN1

**Extended Data Fig. 4 | Modeling and hydrogen-deuterium exchange analysis of 0152-induced stabilization at the APP-CAPRIN1 interface.**

a, Structural model of APP-CAPRIN1-0152 complex generated by docking APP intracellular domain to in silico-modeled CAPRIN1 using HDOCK; 0152 positioned in the drug-binding pocket.

b, HDX-MS protection (blue) and deprotection (pink) mapped onto the APP-CAPRIN1 complex, showing interactions.

c, Regions of reduced deuterium uptake with 0152 indicating stabilized binding clusters on APP intracellular domain.

d, Close-up of APP (cyan) and CAPRIN1 (magenta) helix–helix interface.

e, Predicted hydrogen bonds between APP K650/K651 and CAPRIN1 residues S179, P176, T103.

f, Model of 0152 docked at CAPRIN1-APP interface near KKK motif, interacting with CAPRIN1 residues D90 and K98.

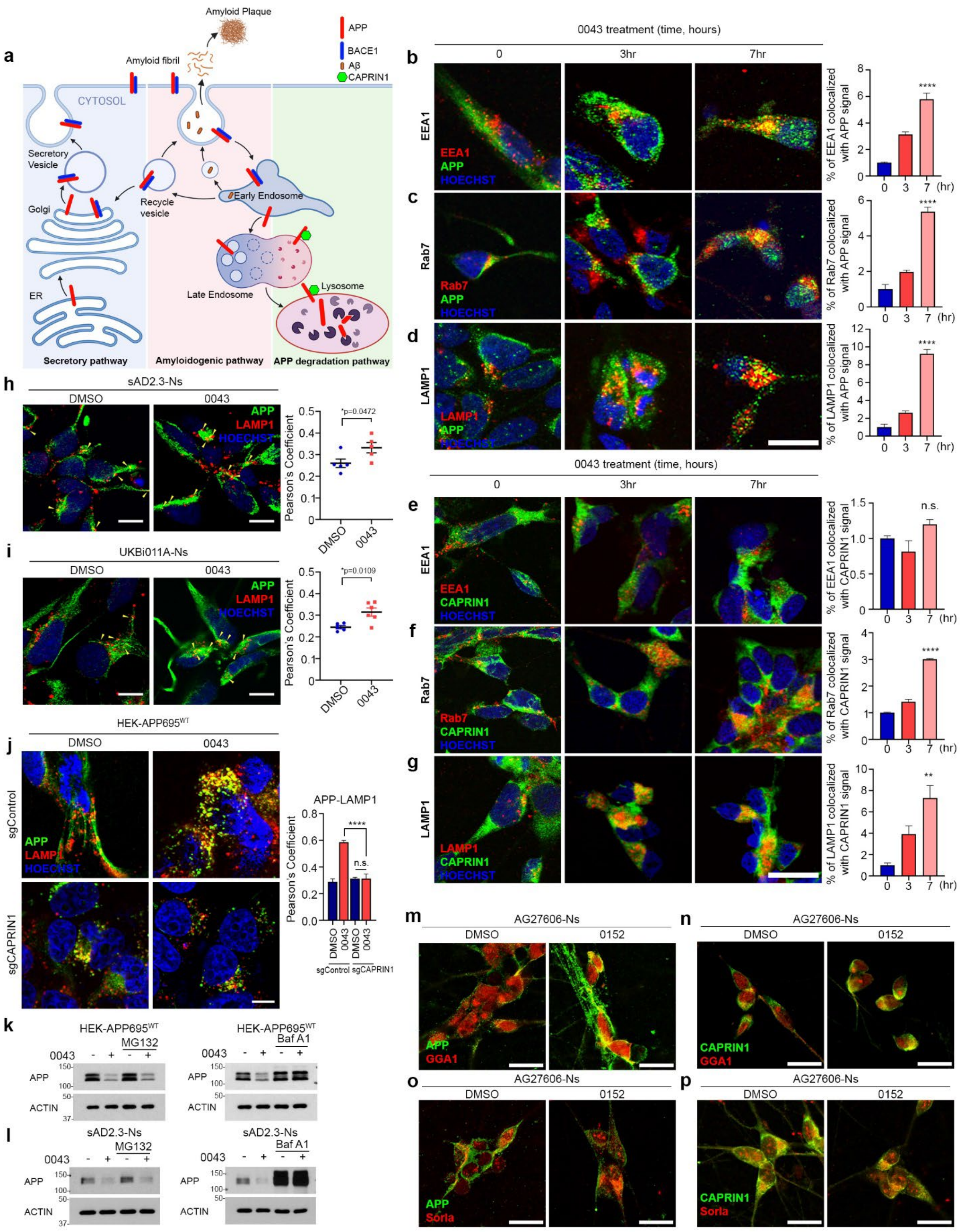

a
Amyloid Plaque
APP
BACE1
Aβ
CAPRIN1
Amyloid fibril
CYTOSOL
Secretory Vesicle
Golgi
ER
Recycle vesicle
Early Endosome
Late Endosome
Lysosome
Secretory pathway
Amyloidogenic pathway
APP degradation pathway
0043 treatment (time, hours)
0
3hr
7hr
b
EEA1
APP
HOECHST
% of EEA1 colocalized with APP signal
c
Rab7
% of Rab7 colocalized with APP signal
d
LAMP1
% of LAMP1 colocalized with APP signal
e
CAPRIN1
% of EEA1 colocalized with CAPRIN1 signal
n.s.
f
% of Rab7 colocalized with CAPRIN1 signal
g
% of LAMP1 colocalized with CAPRIN1 signal
(hr)
h
sAD2.3-Ns
DMSO
0043
*p=0.0472
Pearson's Coefficient
i
UKBi011A-Ns
*p=0.0109
j
HEK-APP695WT
sgControl
sgCAPRIN1
APP-LAMP1
k
MG132
Baf A1
ACTIN
l
m
AG27606-Ns
0152
GGA1
n
o
Sorla
p

**Extended Data Fig. 5 | Small-molecule glue degraders enhance CAPRIN1-dependent endolysosomal trafficking and degradation of APP.**

a, Schematic of APP amyloidogenic and degradation pathways, highlighting CAPRIN1-mediated lysosomal degradation.

b–i, Time-course confocal images of APP (b–d) and CAPRIN1 (e–g) trafficking in AG27606-Ns treated with 0043 (10 µM) for 0, 3, or 7 h; co-stained with EEA1 (b,e), Rab7 (c,f), or LAMP1 (d,g) and Hoechst. Quantification of colocalization shown (mean ± s.d.; one-way ANOVA; n.s., $^{**}p < 0.01$, $^{****}p < 0.0001$); scale bars, 20 µm.

h,i, APP-LAMP1 colocalization in sAD2.3 (h) and UKB011A neurons (i) treated with DMSO or 0043 (10 µM, 24 h) (mean ± s.d.; $^{*}p < 0.05$); scale bars, 20 µm.

j, Confocal images of sgControl and sgCAPRIN1 HEK-APP695$^{WT}$ cells treated with DMSO or 0043 (10 µM, 24 h) stained for APP, LAMP1, and Hoechst (scale bar, 10 µm); APP-LAMP1 colocalization quantified (mean ± s.d., n = 3; n.s., $^{****}p < 0.0001$).

k,l, Western blots of APP in HEK-APP695$^{WT}$ (k) or sAD2.3-Ns (l) pretreated with MG132 or Bafilomycin A1, followed by 0043 (10 µM, 24 h).

m,n, Confocal images of AG27606-Ns treated with DMSO or 0152 (10 µM, 24 h) stained for APP/GGA1 (m) or CAPRIN1/GGA1 (n); scale bars, 20 µm.

o,p, Confocal images of AG27606-Ns treated with DMSO or 0152 stained for APP/Sorla (o) or CAPRIN1/Sorla (p); scale bars, 20 µm.

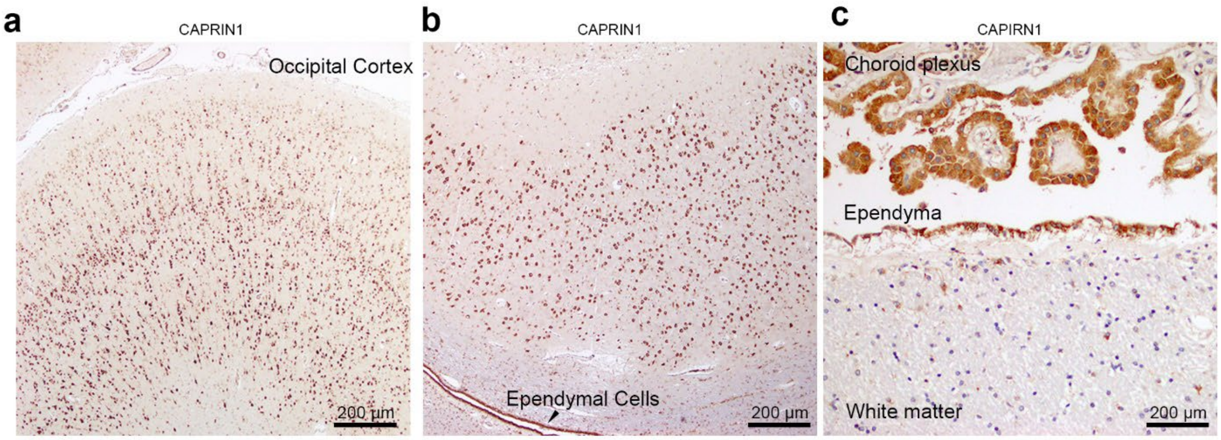


d

| Samples | #1 | #2 | #3 | #4 | none |
|---|---|---|---|---|---|
| Age, Sex | 70, F | 64, F | 67, M | 67, F | 64, F |
| Path Diagnose | AD | AD | AD | AD | No |
| Neuron | ++++ | ++++ | ++++ | ++++ | ++++ |
| Oligodendroglia | - | - | - | - | - |
| astroglia | +/- | +/- | +/- | +/- | +/- |
| ependymal cells | +++ | +++ | +++ | +++ | +++ |
| Choroid Plexus | +++ | +++ | +++ | +++ | +++ |
| Endothelial cells | - | - | - | - | - |

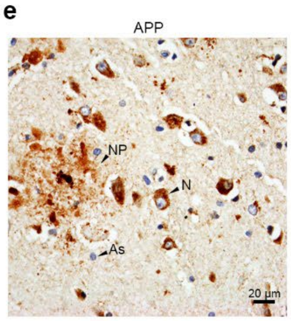


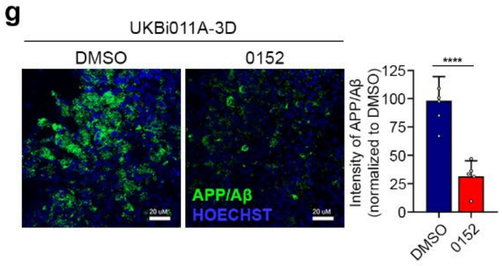


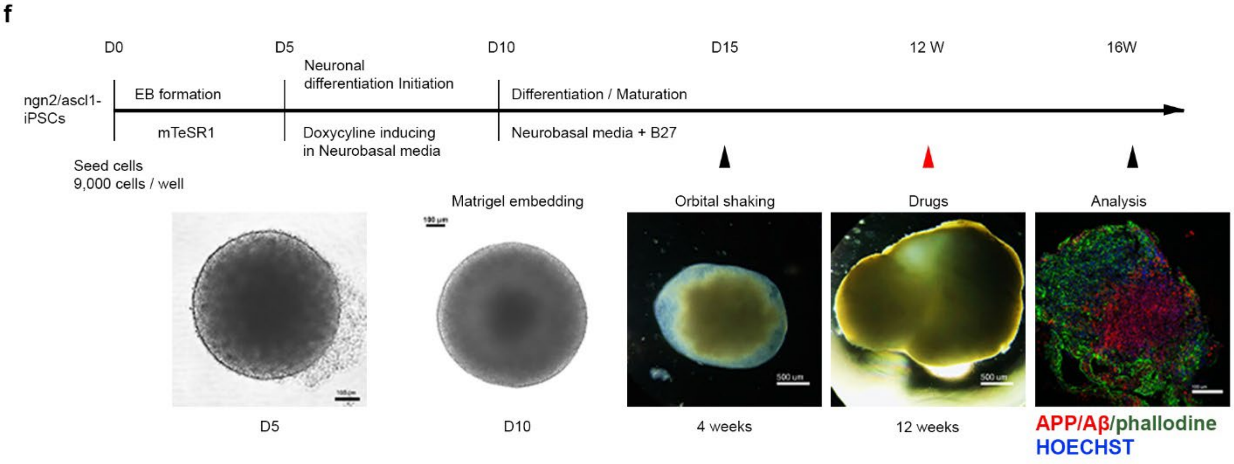

**Extended Data Fig. 6 | CAPRIN1 expression in human AD brains and 0152-mediated APP/Aβ reduction in 3D cortical organoids.**

a–c, IHC of CAPRIN1 in AD patient brain sections showing expression in occipital cortex (a), ependymal cells lining ventricles (b), and choroid plexus (c); scale bars, 200 μm.

d, Summary of pathological classification of AD and non-AD patient brain samples.

e, APP/Aβ IHC in cortex showing neuronal (N), astrocytic (As) and neuritic plaque (NP) labeling (scale bar, 20 μm).

f, Timeline and representative images of brain organoid generation from Ngn2/Ascl1-iPSCs: EB aggregation (5 days), doxycycline-induced neuronal differentiation (5 days), Matrigel embedding at D10, orbital shaking from D15, drug treatment at week 12, and representative organoid at week 16 stained for APP/Aβ (red), phalloidin (green), Hoechst (blue).

g, Confocal images and quantification of APP/Aβ intensity in UKBi011A-derived 3D organoids treated with DMSO or 0152 (10 μM) (scale bar, 20 μm; mean ± s.d.; $n \geq 4$).

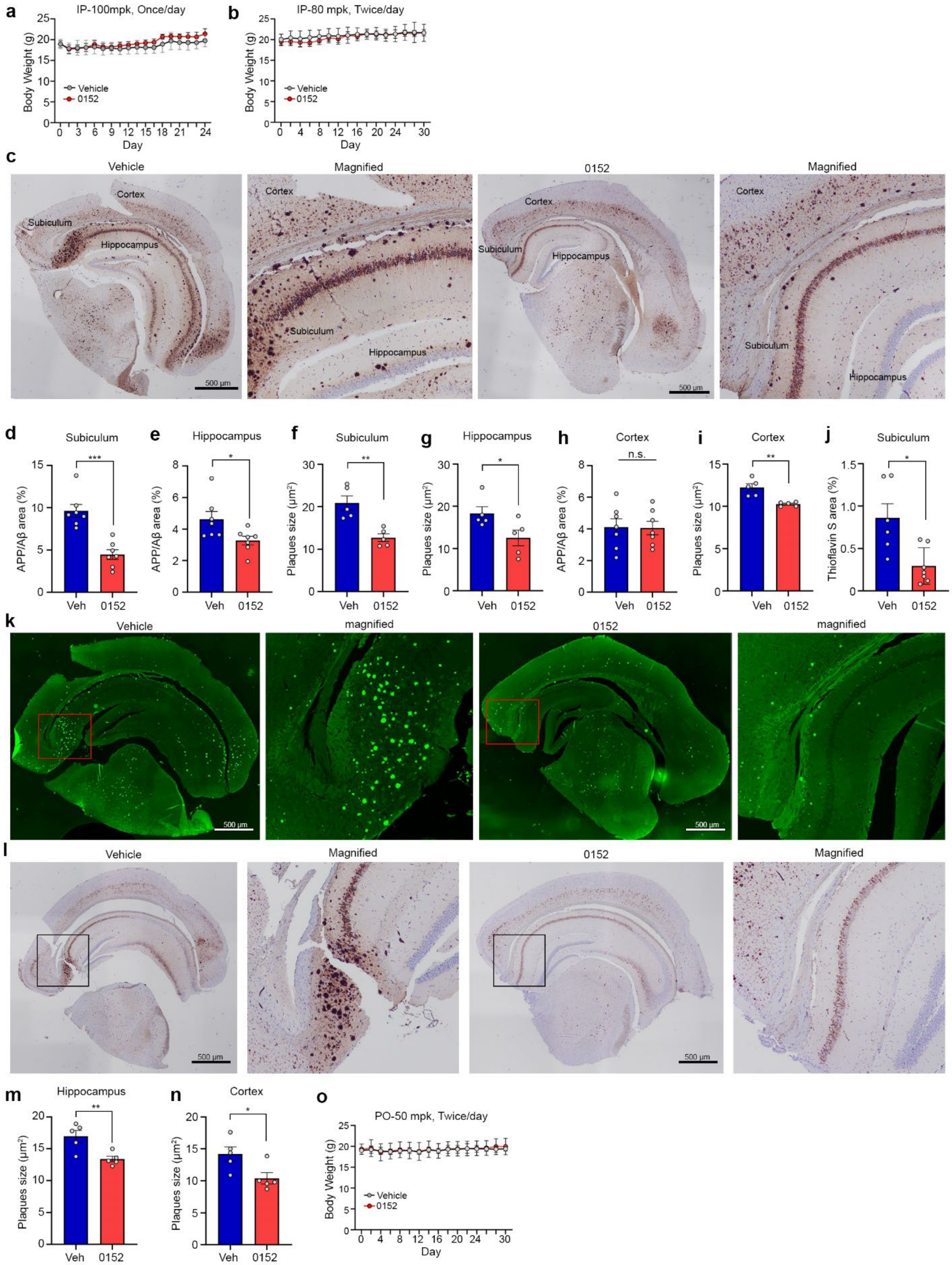
a
IP-100mpk, Once/day
Body Weight (g)
Day
Vehicle
0152
b
IP-80 mpk, Twice/day
Body Weight (g)
Day
Vehicle
0152
c
Vehicle
Magnified
0152
Magnified
Cortex
Subiculum
Hippocampus
500 μm
d
Subiculum
APP/Aβ area (%)
***
Veh
0152
e
Hippocampus
APP/Aβ area (%)
*
f
Subiculum
Plaques size (μm²)
**
g
Hippocampus
Plaques size (μm²)
*
h
Cortex
APP/Aβ area (%)
n.s.
i
Cortex
Plaques size (μm²)
**
j
Subiculum
Thioflavin S area (%)
*
k
Vehicle
magnified
0152
magnified
500 μm
l
Vehicle
Magnified
0152
Magnified
500 μm
m
Hippocampus
Plaques size (μm²)
**
Veh
0152
n
Cortex
Plaques size (μm²)
*
Veh
0152
o
PO-50 mpk, Twice/day
Body Weight (g)
Day
Vehicle
0152

**Extended Data Fig. 7 | Therapeutic effects of 0152 following IP and oral administration in 5xFAD mice.**

a,b, Body weights during 0152 treatment at different dosing regimens: (a) IP once daily 100 mg/kg; (b) IP twice daily 80 mg/kg (mean ± s.e.m.).

c, IHC of 5xFAD mouse brains showing APP/Aβ accumulation in cortex, hippocampus, and subiculum after 30-day vehicle or 0152 treatment (80 mg/kg, IP twice); scale bars, 500 μm.

d–i, Quantification of APP/Aβ-positive area and plaque size in subiculum, hippocampus, and cortex (mean ± s.e.m.; $*p < 0.05$, $**p < 0.01$, $***p < 0.001$; n.s.; unpaired two-tailed t-test).

j,k, Thioflavin S staining (j) and quantification of Thioflavin S-positive area in subiculum (k) after 0152 treatment (mean ± s.e.m.; $*p < 0.05$; unpaired two-tailed t-test).

l, IHC showing APP/Aβ accumulation after 30-day vehicle or 0152 treatment by oral gavage (80 mg/kg, PO twice); scale bars, 500 μm.

m,n, Quantification of APP/Aβ plaque size in hippocampus (m) and cortex (n) after oral treatment (mean ± s.e.m.; $*p < 0.05$, $**p < 0.01$).

o, Body weight monitoring during PO twice daily 80 mg/kg 0152 treatment (mean ± s.e.m.).

**Supplementary Table 1. Oligonucleotides**

| | | |
|---|---|---|
| **CRISPR** | | |
| Caprin1 (Oligo1) | IDT | 5'-CACCGCGACAAGAAACTTCGGAACC-3' |
| Caprin1 (Oligo2) | IDT | 5'AAACGGTTCCGAAGTTTCTTGTCG C-3' |
| Control primer | Addgene | 80248 |
| **qPCR primers** | | |
| APP (Forward) | IDT | 5′-TGACAAGTTCCGAGGGGTAG-3′ |
| APP (Reverse) | IDT | 5′-CCAGACATCCGAGTCATCCT-3′ |
| GAPDH (Forward) | IDT | 5′-CTCTGACTTCAACAGCAC-3′ |
| GAPDH (Reverse) | IDT | 5′-CATACCAGGAAATGAGCTTGACAA-3′ |

#### Referernce for Methods
